# A novel non-catalytic scaffolding activity of Hexokinase 2 contributes to EMT and metastasis

**DOI:** 10.1101/2021.04.08.439049

**Authors:** Catherine Blaha, Gopalakrishnan Ramakrishnan, Sang-Min Jeon, Veronique Nogueira, Hyunsoo Rho, Soeun Kang, Prashanth Bhaskar, Alexander R. Terry, Alexandre F. Aissa, Maxim V. Frolov, Krushna C. Patra, R. Brooks Robey, Nissim Hay

## Abstract

Hexokinase 2 (HK2), a glycolytic enzyme that catalyzes the first committed step in glucose metabolism, is markedly induced in cancer cells. HK2’s role in tumorigenesis has been attributed to its glucose kinase activity. However, we uncovered a novel kinase-independent HK2 activity, which promotes metastasis. We found that HK2 binds and sequesters glycogen kinase 3 (GSK3) and acts as a scaffold forming a ternary complex with the regulatory subunit of protein kinase A (PRKAR1a) and GSK3b to facilitate GSK3b phosphorylation by PKA, and to inhibit its activity. Thus, HK2 functions as an A-kinase anchoring protein (AKAP). GSK3b is known to phosphorylate proteins, which in turn are targeted for degradation. Consistently, HK2 increased the level and stability of the GSK3 targets, MCL1, NRF2, and SNAIL. In a mouse model of breast cancer metastasis, systemic HK2 deletion after tumor onset inhibited metastasis, which is determined by the effect of HK2 on GSK3b and SNAIL. We concluded that HK2 promotes SNAIL stability and breast cancer metastasis via two mechanisms: direct modulation of GSK3-activity and SNAIL- glycosylation that decreases susceptibility to phosphorylation by GSK3.

---

Increased glucose uptake and intracellular glucose metabolism are known to occur in cancer cells in response to hyperactivation of signaling pathways ^1^. However, little is known about retrograde signaling, whereby glycolytic enzymes regulate the activity of signaling enzymes. The first committed step in glucose metabolism is catalyzed by hexokinases. The phosphorylation of glucose by hexokinases determines the flux of glucose in not only in glycolysis but also in the pentose phosphate pathway (PPP) and the hexosamine biosynthetic pathway ^2^. Among the hexokinase isoforms hexokinase 1 (HK1) and hexokinase 2 (HK2) are mitochondria-associated, high affinity, low-Km hexokinases that couple oxidative phosphorylation and glycolysis. While HK1 is widely expressed in mammalian tissues, HK2 is expressed only in a limited number of tissues. However, when normal cells are converted to cancer cells, they start expressing very high levels of HK2 in addition to the already expressed HK1. Therefore, HK2 expression is considered a hallmark of cancer cells that determines their high glycolytic flux. Thus far, the role of HK2 in tumorigenesis was attributed to its glycolytic activity ^3, 4^.

The low-Km hexokinases (HK1 and HK2) are inhibited by their own catalytic product, glucose-6-phosphate (G6P), which distinguishes them from glucokinase. G6P acts as an allosteric inhibitor of HK1 and HK2 and this feedback inhibition is likely instated to coordinate the uptake of glucose and ATP consumed to phosphorylate it with the cellular demand for glucose as a carbon source for energy and building blocks for anabolic processes. Notably, G6P mimetics could be used to target HK2 glycolytic activity for cancer therapy ^4, 5^. Buildup in G6P level may occur in a temporal manner or when the flux downstream of G6P is attenuated or inhibited. The substitution of 2-deoxyglucose (2-DG) for glucose results in 2-deoxyglucose-6-phosphate (2- DG6P) accumulation because 2-DG6P cannot be further metabolized in glycolysis. Although 2- DG6P is less efficient than G6P in inhibiting HK1 and HK2, its rapid accumulation to relatively high amounts inside cells inhibits HK1 or HK2 ^6, 7^. Thus, 2-DG can be used to study the consequences of the allosteric inhibition of HK1 and HK2 by their own catalytic product. Using this approach, we uncovered a novel mechanism by which HK2 affects the activity of glycogen kinase 3 (GSK3).

GSK3 is a Ser/Thr kinase, which plays a crucial role in many vital cellular processes, such as cell proliferation, apoptosis, metabolism, and cancer progression. There are two isoforms of GSK3 (*α* and *β*) encoded by two separate genes. The kinase activity of GSK3 is inhibited by phosphorylation of the protein at Ser21/Ser9 (corresponding residues in GSK3 α/β). The phosphorylation of GSK3 on these residues is mediated by several AGC kinases ^8–12^, and by Ser/Thr phosphatases such as protein phosphatase 1 (PP1) and protein phosphatase 2A (PP2A) ^13^.

Our results showed that HK2 sequesters GSK3 inhibiting its activity and accessibility to its targets. Importantly, we demonstrated that HK2 directly binds GSK3b and the cyclic-AMP dependent protein kinase A (PKA) regulatory subunit 1a (PRKAR1a) to facilitate GSK3b phosphorylation by PKA. Thus, HK2 functions as AKAP, independently of its catalytic activity. However, G6P disrupts the binding of HK2 to GSK3b and PRKAR1a. In vivo, the accumulation of G6P or 2DG6P, which change allosterically HK2, dramatically reduced the inhibitory phosphorylation of GSK3*α* and GSK3*β* on Ser21 or Ser9 respectively, by dissociating GSK3 and increasing its susceptibility to PP2A.

Phosphorylation of proteins by GSK3 often targets them for degradation ^14^. Consistent with the effect of HK2 on GSK3 activity, we found that both wild-type (WT) HK2 and kinase-dead HK2 affect the level and stability of MCL-1, NRF2, and SNAIL which are known GSK3 targets ^15–17, 18^. We have demonstrated that SNAIL’s stabilization by HK2 promotes EMT and breast cancer metastasis.

## Results

### The glucose analog 2-deoxyglucose (2-DG) inhibits the phosphorylation of glycogen synthase kinase 3 (GSK3) and increases its activity

The replacement of glucose with two different glucose analogs, 2-DG and 5-thioglucose (5-TG), revealed a rapid reduction in the phosphorylation of GSK3*β* specifically by 2-DG, and not by 5- TG, in both Rat1a cells and mouse embryo fibroblasts (MEFs) (Fig. 1a). Although neither 2-DG nor 5-TG can be metabolized in glycolysis, 2-DG can be phosphorylated to 2-DG6P by hexokinase, whereas 5-TG, although binds hexokinase, cannot be phosphorylated by hexokinase (Fig. 1b). The effect of 2-DG on both GSK3*α* and GSK3*β* phosphorylation was observed in most rodent and human cell lines that were tested (Fig. 1c).

**Figure 1:**
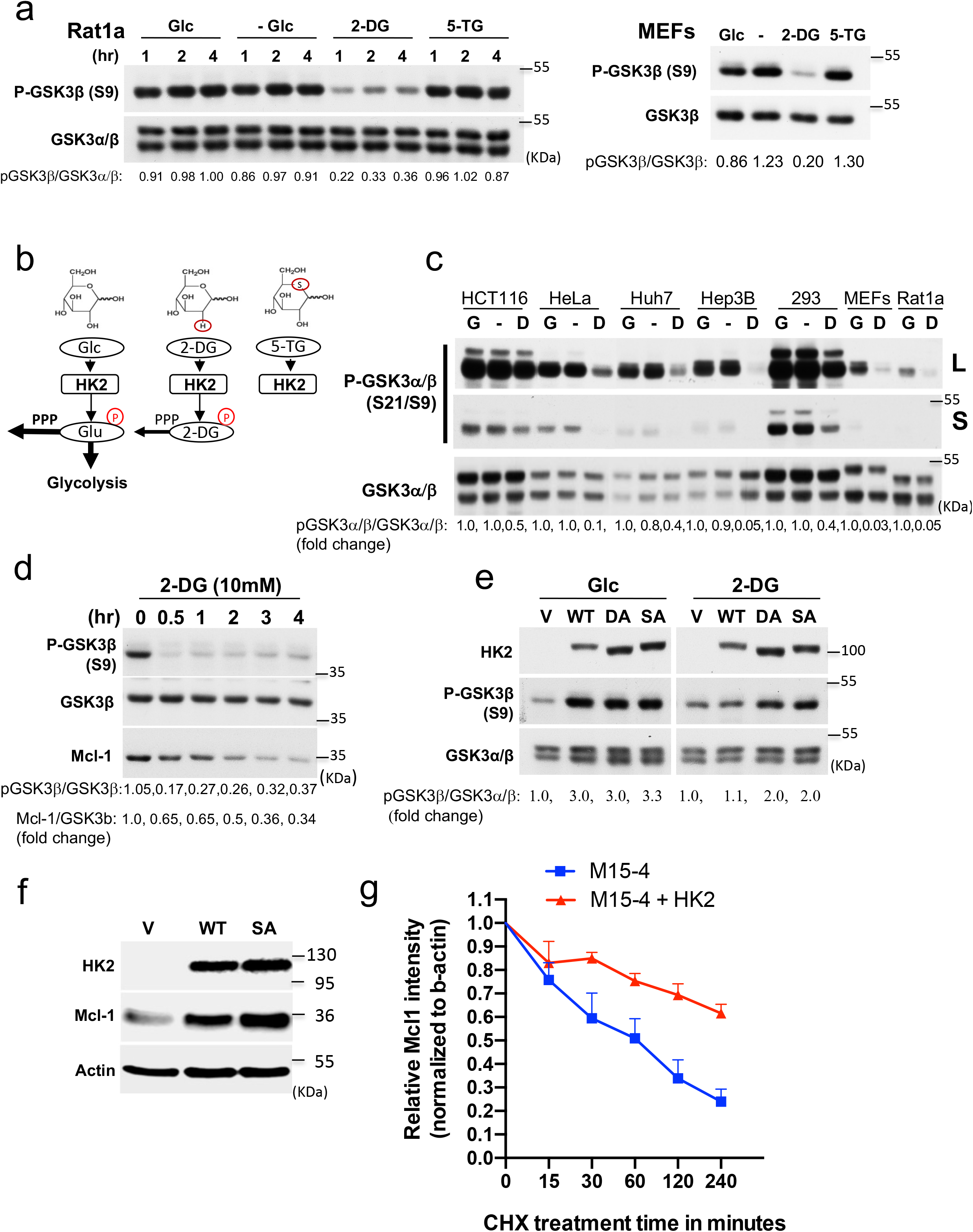
Replacement of glucose with 2-DG inhibits the phosphorylation of GSK3 and increases GSK3 activity in a HK2-dependent manner. **a.** Rat1a cells or MEFs were incubated in glucose free medium in the absence (-) or presence of 10mM glucose (Glc), 2-DG or 5-TG. Immunoblots showing GSK3*β* phosphorylation, at the indicated time points after incubation. **b.** Schematic depicting the structures of glucose (Glc), 2-DG, and 5-TG and their utilization by HK inside cells. Similar to Glc, 2-DG can be phosphorylated by HK2 but cannot be further metabolized except in the first step of the pentose phosphate pathway (PPP). 5-TG cannot be phosphorylated by HK2 **c.** Cells were incubated in the presence of glucose (G), absence of glucose (-) or presence of 2-DG (D) for 2 hr. An immunoblot image shows the phosphorylation of GSK3*α* and GSK3*β*. **d.** Rat1a cells were incubated in glucose-free medium in the presence of 10 mM 2-DG for the indicated durations. Cells were then harvested for immunoblotting to determine GSK3*β*phosphorylation and MCL1 levels. **e.** MI5-4 CHO cells expressing either wild-type (WT) HK2, individual kinase-dead HK2 mutants (DA, SA) or empty vector (V) were incubated in glucose-free medium in the presence of 10 mM glucose (G) or 2-DG (D). After 2 hr, cells were harvested and analyzed for immunoblotting. **f.** The level of the Mcl-1 protein after expression of WT or kinase-dead SA mutant HK2 in M15- 4 CHO cells. **g.** Protein stability of MCL-1 in M15-4 CHO cells and M15-4 CHO cells expressing WT HK2 as measured after exposure to cycloheximide (CHX). Plot showing MCL1 protein half-life after quantification relative to b-actin in 3 independent experiments.

GSK3 is a versatile kinase that participates in many fundamental cellular processes, such as cell proliferation, cell growth, and cell survival ^9, 19^. In many cases the phosphorylation of proteins by GSK3 targets them for degradation. For instance, GSK3 was shown to regulate the stability of the anti-apoptotic protein Mcl-1 through direct phosphorylation ^15^. Indeed, we found that replacement of glucose with 2-DG markedly and rapidly reduced the level of Mcl-1 (Fig. 1d).

### HK2 maintains a high level of GSK3 phosphorylation independent of its activity, but its activity is required for the suppression of GSK3 phosphorylation by 2-DG

Unlike 5-TG, 2-DG can be phosphorylated by hexokinase, so we examined whether hexokinase activity is required for the inhibitory effect of 2-DG on GSK3*β* phosphorylation. We took advantage of M15-4 CHO cells ^20^, which lack detectable hexokinase expression and have limited hexokinase activity, and ectopically expressed either WT HK2 or HK2 mutants in these cells. HK2- DA is a mutant in which alanine is substituted for two aspartic acid residues in the amino- and carboxy-terminal domains of HK2 required for the binding of glucose (D209A/D657A). HK2-SA is a mutant in which alanine is substituted for two serine residues in the amino- and carboxy-terminal domains required for catalytic activity (S155A/S603A) ^21^. HK2-dMT is a mutant that carries a deletion of the first 20 amino acids, which are required for binding to VDAC and mitochondria. When expressed in M15-4 CHO cells, both WT HK2 and dMT mutant of HK2 had catalytic activity, whereas the DA and SA mutants had very little or no catalytic activity (Extended Data Fig. 1A). As shown in Fig. 1e and Extended Data Fig. 1b, 2-DG had no effect on GSK3*β* phosphorylation in M15-4 CHO cells expressing vector alone. However, expression of WT HK2 in these cells elevated GSK3*β* phosphorylation in the presence of glucose, but GSK3*β*phosphorylation was decreased in the presence of 2-DG. Surprisingly, both WT HK2 and kinase-inactive HK2 mutants increased GSK3*β* phosphorylation in the presence of glucose (Fig. 1e). However, the replacement of glucose with 2-DG reduced GSK3*β* phosphorylation in cells expressing kinase-active WT HK2 or the mitochondrial binding-deficient mutant (dMT-HK2) and, to a much lesser extent, in cells expressing kinase-inactive mutants (Fig. 1e, and Extended Data Fig. 1b). Interestingly, overexpression of WT HK2, kinase-inactive HK2 mutants, or a mitochondrial binding-deficient mutant of HK2 even in cells that express endogenous hexokinases, such as Rat1a and HEK-293 cells, also increased GSK3*β* phosphorylation (Extended Data Fig. 2a, b). However, in all cell lines tested, 2-DG markedly reduced GSK3*β* phosphorylation only in cells expressing kinase-active HK2 (Fig. 1e, and Extended Data Fig. 2a, b). Taken together, these results indicate that high-level expression of HK2 can elevate the phosphorylation of GSK3*β* in a kinase-independent manner, while 2-DG markedly inhibits GSK3*β* phosphorylation only in the presence of a catalytically active hexokinase. The ability of HK2 to affect GSK3*β* phosphorylation appears to be independent of its binding to mitochondria. Therefore, these data strongly suggest that the effect of 2-DG on GSK3 phosphorylation is dependent upon 2-DG6P. Moreover, this can explain why 5-TG had no effect on the phosphorylation of GSK3, as unlike 2- DG, 5-TG cannot be phosphorylated. (see Fig. 1a, b). Consistent with the effect of HK2 on GSK3 phosphorylation and activity, we found that the expression of either WT or mutant HK2 in M15- 4 CHO cells markedly increased the steady state levels and stability of Mcl-1 (Fig. 1f and Fig. 1g).

### The effects of HK2 and 2-DG on GSK3β phosphorylation are mediated by PKA and PP2A

The phosphorylation of GSK3*β*on Ser9, leads to its inhibition and is mediated by several AGC kinases including AKT, PKC, PKA, p70S6K, and p90RSK as well as Ser/Thr protein phosphatases such as PP1 and PP2A ^22, 23^. To understand the mechanism by which GSK3 phosphorylation is affected, we first determined which of the known kinases that phosphorylate GSK3*β* on Ser9 might be involved. Akt is usually the predominant kinase that phosphorylates GSK3. However, we excluded Akt involvement because 2-DG inhibited GSK3*β*phosphorylation independently of Akt activity (Extended Data Fig. 3a-c). First, 2-DG reduced GSK3*β* phosphorylation despite a corresponding increase in Akt activation. Second, although the complete depletion of Akt activity in Akt1/2 DKO MEFs by LY294002 (PI3-kinase inhibitor) treatment reduced GSK3*β*phosphorylation in the presence of glucose, it was further reduced when glucose was replaced with 2-DG. Finally, overexpression of a constitutively activated form of Akt (mAkt) was associated with GSK3*β* phosphorylation, which was largely amenable to 2-DG inhibition. Taken together, these findings suggest that the effects of 2-DG on GSK3*β*phosphorylation and activation are not dependent on Akt.

In an unbiased yeast two-hybrid screen, where a full length HK2 was used as a bait, we found that a protein fragment containing the first 129 amino acids, which encompasses the dimerization/docking domain of PRKAR1a ^24^, the regulatory subunit of PKA, interacts with HK2 (Extended Data Fig. 4). Therefore, we next explored the possibility that the effect of 2-DG6P on GSK3*β* phosphorylation is mediated by PKA. PRKAR1a (RIa) binds the catalytic subunit of PKA and restrains its activity, but upon binding of cyclic-AMP, the catalytic subunit is dissociated from PRKAR1a, enabling the phosphorylation of PKA target proteins ^25^. We, therefore, focused on the possibility that the effect of HK2 on GSK3 phosphorylation is mediated by PKA. We examined the effect of the adenylate cyclase inhibitor, 2’5’-dideoxyadenosine on the ability of HK2 to increase the phosphorylation of GSK3. As shown in Fig. 2a, the increase in GSK3 phosphorylation following overexpression of HK2 in M15-4 CHO cells was blunted in the presence of the adenylate cyclase inhibitor 2’5’-dideoxyadenosine. We then used the PKA inhibitor H-89 and found that it reduced GSK3 phosphorylation mediated by HK2 (Fig. 2b). To further understand the mechanism by which HK2 affects GSK3 phosphorylation through PRKAR1a, we used immunoprecipitation experiments with HK2 and PRKAR1a to confirm the yeast two-hybrid results. Indeed, we found that HK2 bound PRKAR1a in the presence of glucose, but this binding was inhibited when glucose was replaced with 2-DG (Fig. 2c). As shown in Fig. 2d and Extended Data Fig. 5a, ectopically expressed HK2 also pulled down ectopically expressed GSK3*β* in the presence of glucose but not in the presence of 2-DG. Either the WT or active GSK3*β* mutant bound HK2 in the presence of glucose, but to a much lesser extent in the presence of 2-DG (Extended Data Fig. 5a). In a reciprocal experiment, GSK3*β* pulled down HK2 (extended Fig. 5b). Notably, GSK3*β* was phosphorylated when was bound to HK2 (Extended Data Fig. 5a). Finally, when exogenous HK2 was immunoprecipitated, it pulled down endogenous PRKAR1a, but only in the presence of glucose and not in the presence of 2-DG (Fig. 2e). Importantly, when M15-4 CHO cells were treated with 2-DG, GSK3*β* was dissociated from only catalytically active HK2 but not from kinase-dead HK2 mutants (Extended Fig. 5c). This is consistent with the results showing that 2-DG markedly decreased GSK3*β* phosphorylation only when kinase-active HK2 was expressed but not when the kinase-dead HK2 mutant was expressed (Fig. 1e and Extended Data Fig. 2b). To verify that endogenous HK2 binds endogenous GSK3*β*, we immunoprecipitated endogenous HK2 from HEK293 cells and found that it interacted with endogenous GSK3*β*, but this interaction did not occur when glucose was replaced with 2-DG (Fig. 2f, see also Extended Data Fig. 13c). Taken together, these results raise the possibility that the binding of PRKAR1a and GSK3*β* to hexokinase increases the susceptibility of GSK3*β* to PKA-mediated phosphorylation in a cyclic AMP- and PKA-dependent manner. However, conformational changes in HK2 in response to 2-DG6P or glucose-6-phosphate (G6P) binding inhibits the interaction between HK2, PRKAR1a, and GSK3*β*and thereby decreases GSK3*β* phosphorylation.

**Figure 2:**
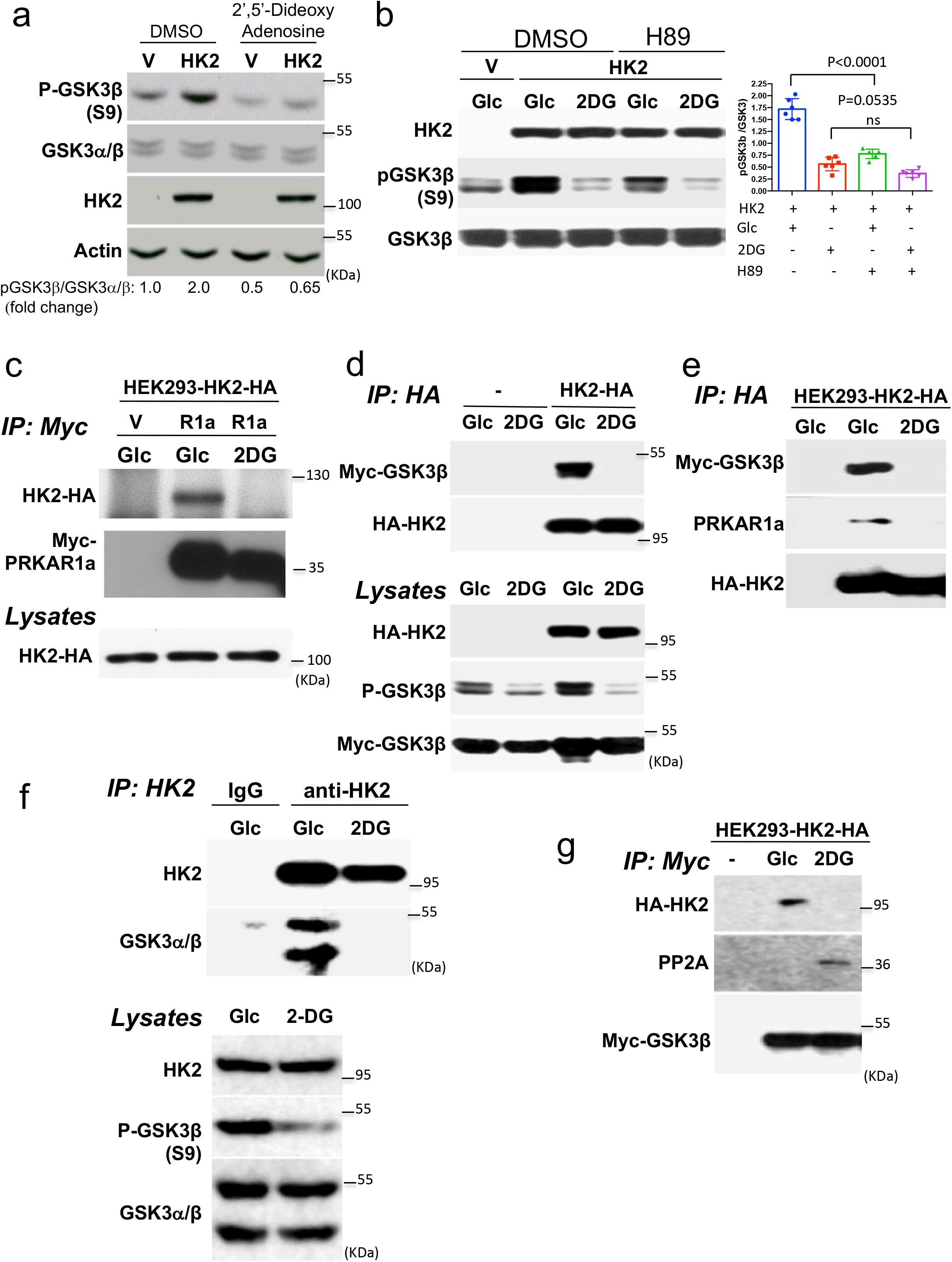
HK2 affects GSK3β phosphorylation in a PKA-dependent manner and interacts with PRKAR1a and GSK3β in a 2-DG-dependent manner. **a.** GSK3*β* phosphorylation following overexpression of HK2 in M15-4 cells in the presence of the adenylate cyclase inhibitor 2’5’-dideoxyadenosine 200uM for 6 hr. **b.** GSK3*β* phosphorylation following overexpression of HK2 in M15-4 cells in the presence of the PKA inhibitor H89. Cells were pretreated with either DMSO or H89 (10uM) for 2hr and then exposed to 10mM glucose (Glc) or 2 DG for another 2hr in the presence of either DMSO or H89. (left panel: representative immunoblot; right panel: quantification of pGSK3*β*/GSK3*β*. Results are the mean ± SEM of 3 independent experiments in duplicates). **c.** After transfection of control myc-tagged vector (V) or myc-tagged PRKAR1a (R1a) or control myc-vector plasmid into HEK293 cells stably expressing HA-tagged HK2 (HEK293-HK2-HA), cells were incubated in glucose-free medium in the presence of 10 mM glucose (Glc) or 2-DG. After 2 hr, cells were lysed for immunoprecipitation with anti-myc antibody, followed by immunoblotting using anti-HA and anti-myc-HRP antibody. Total lysates were subjected to immunoblotting using anti-HA antibody. **d.** Control or HK2-HA stably expressing cells were transfected with myc-GSK3β. Cells were then incubated in glucose-free medium in the presence of 10 mM glucose (Glc) or 2-DG. After 2 hr, cells were lysed for immunoprecipitation with anti-myc antibody, followed by immunoblotting using anti-HA and anti-myc-HRP antibodies. Total lysates were subjected to immunoblotting using anti-p-GSK3β, anti-myc-HRP, and anti-HA antibodies. **e.** After transfection of myc-GSK3β into HEK293-HK2-HA cells, cells were incubated in glucose-free medium in the presence of 10 mM glucose (Glc) or 2-DG. After 2 hr, cells were lysed for immunoprecipitation with anti-HA antibody, followed by immunoblotting using anti-myc-HRP antibody and anti-PRKAR1a antibody. First lane shows control untransfected cells. **f.** HEK293 cells were incubated in glucose-free medium in the presence of 10 mM glucose (Glc) or 2-DG. After 2hr, cells were lysed for immunoprecipitation. Endogenous HK2 was immunoprecipitated with anti-HK2 and subjected to immunoblotting with anti-HK2 and anti- GSK3*α*/*β*. **g.** The experiment was performed as described in D, except that after immunoprecipitation with anti-myc, immunoblotting was performed with anti-PP2A.

Since both the protein phosphatase 1 (PP1) and protein phosphatase 2A (PP2A) phosphatases also regulate GSK3 phosphorylation ^13^, we examined their potential involvement in the effect of hexokinase on GSK3*β* phosphorylation. Treatment of cells with the PP2A inhibitor okadaic acid (OA) or the PP1 inhibitor tautomycetin (TC) revealed that treatment with 100 nM OA markedly increased GSK3*β* phosphorylation in the presence of glucose but also blunted the decrease in GSK3 phosphorylation after glucose was replaced with 2-DG (Extended Data Fig. 6a). TC failed to mimic these effects at concentration as high as 500nM (Extended Data Fig. 6a), but the specific PP2A inhibitor LB100 inhibited the effect of 2-DG on GSK3*β* phosphorylation (Extended Data Fig. 6b), suggesting PP2A involvement in the dephosphorylating effect of 2-DG on GSK3*β*. Endogenous PP2A interaction with GSK3*β* dissociated from HK2 in the presence of 2- DG (Fig. 2g) is compatible with this interpretation. These results indicate that HK2 facilitates GSK3 inhibition not only via PKA-mediated phosphorylation, but also via physical GSK3 sequestration and the associated prevention of its dephosphorylation and activation by PP2A.

### HK2 forms complexes with GSK3b and PRKAR1a (RIa) in vitro in a G6P-dependent manner, indicating direct binding

As was shown for bone-fide AKAPs the ultimate definition of AKAP is determined by its ability to bind directly to the PKA regulatory subunits together with the target protein. To further determine if HK2 can be categorized as AKAP, we conducted in vitro binding assays as previously described ^26^. We employed His-tagged or GST-tagged protein coated plates. The plates were subjected to the interacting proteins and interaction was determined by fluorescent anti-tagged antibodies against the interacting protein (Fig. 3, Left panels). As shown in Fig. 3a anchored Ria binds the increasing amounts of HK2, but this interaction was disrupted after addition of G6P. Likewise, anchored GSK3b binds HK2, and this binding is disrupted by the addition of G6P (Fig. 3b). Finally, anchored RIa binds GSK3b only when HK2 is present to form a ternary complex (Fig. 3c). This complex is also sensitive to the addition of G6P but to a less extent than the individual complexes suggesting that the ternary complex is more resistant to the allosteric inhibition of HK2. To determine if the ternary complex can bind the catalytic PKA subunit (PKAc), we added PKAc to the tethered complex and showed that it binds in an HK2 and RIa dependent manner (Fig. 3d). Furthermore, addition of cAMP markedly induced the phosphorylation of GSK3b in the complex, which was diminished by G6P (Fig. 3e). Therefore, collectively, these results provide a strong evidence that the binding of HK2 to GSK3b and RIa is direct and that HK2 is a bone-fide AKAP.

**Figure 3.**
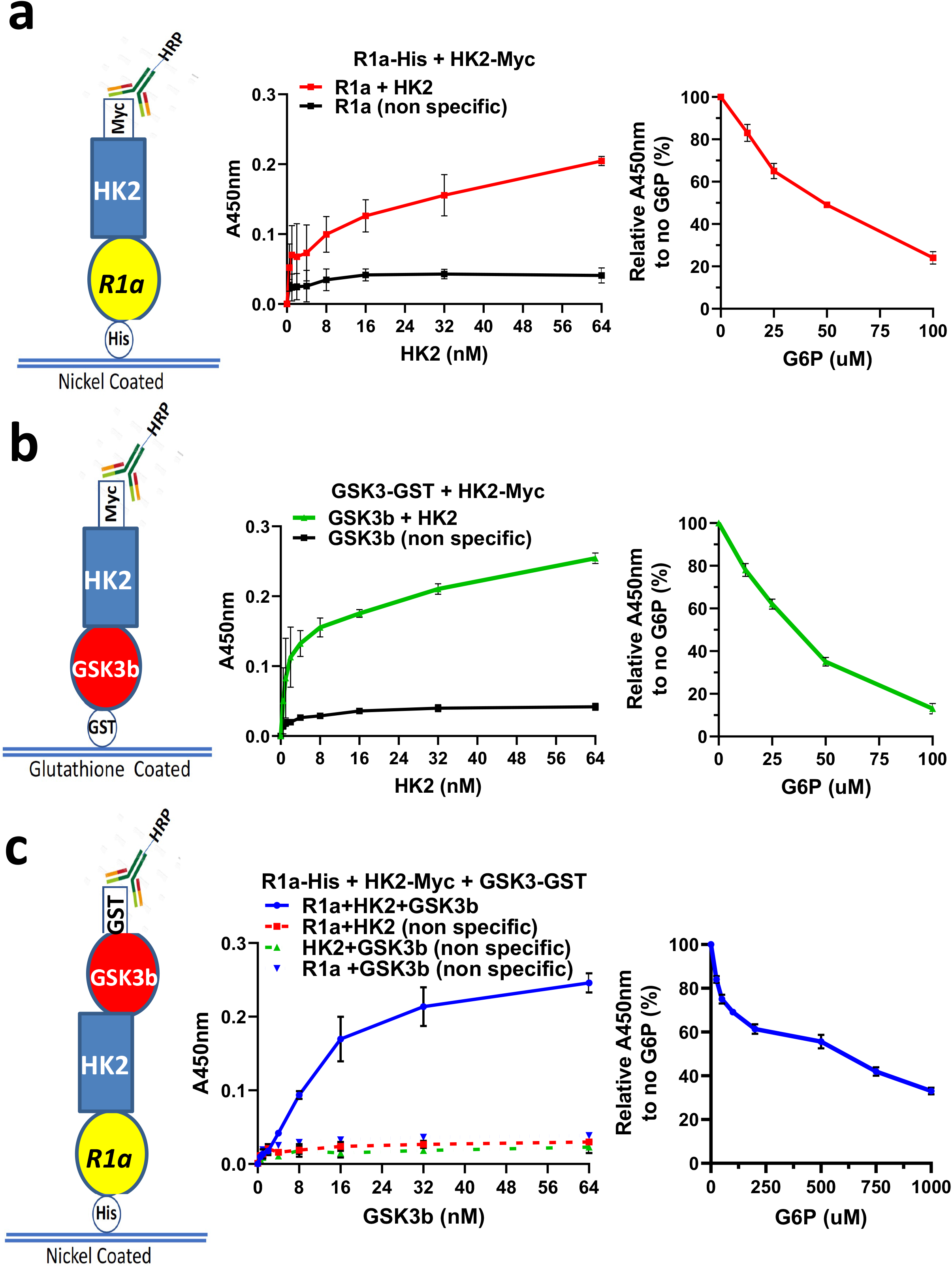
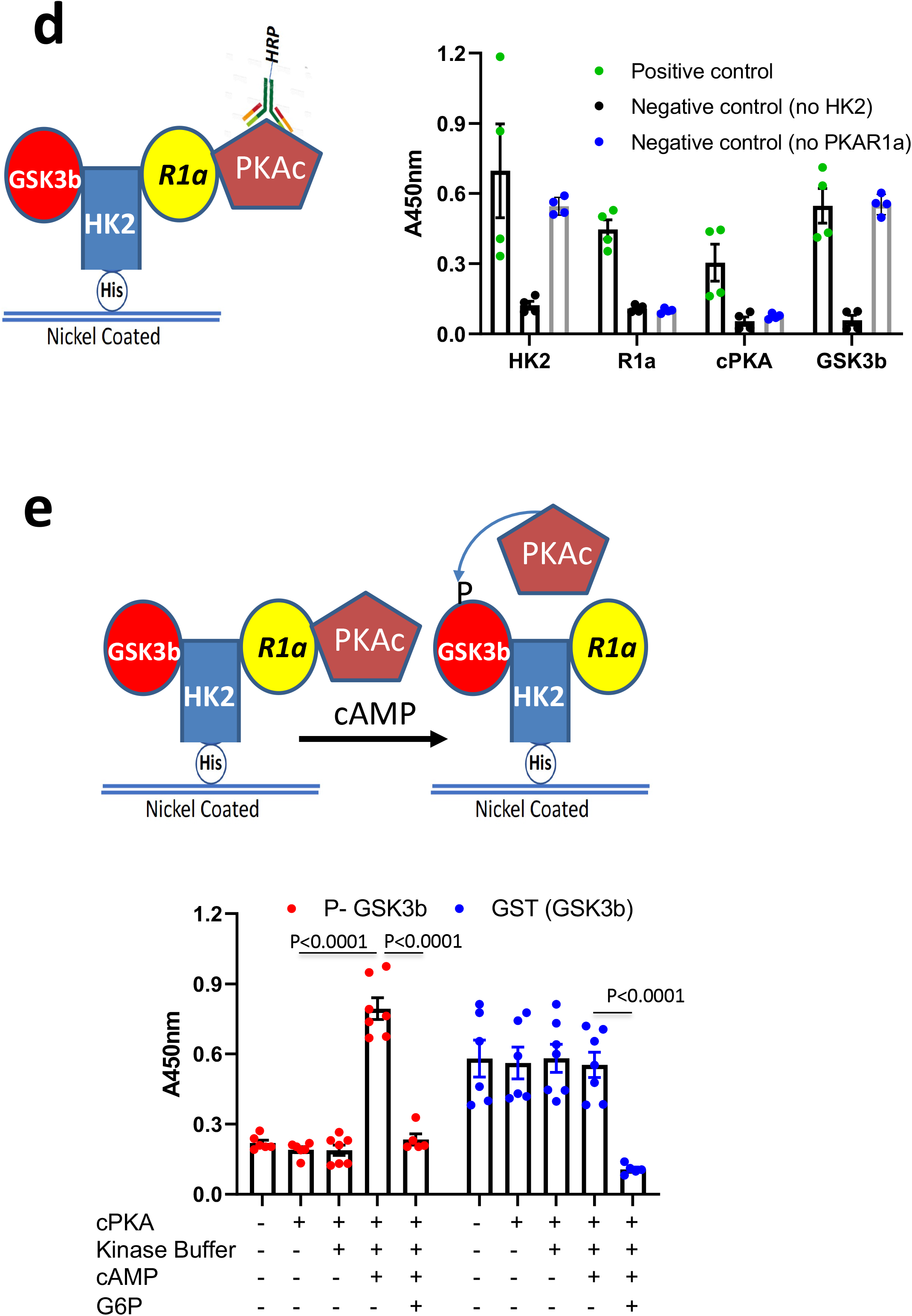
HK2 directly binds GSK3b and R1a in vitro to form complexes that are disrupted by G6P. **a.** Nickel-coated 96-well plates were incubated overnight at 4°C with His-PRKAR1a (50 nM) and then incubated with Myc-HK2 (0.25– 64 nM) in blocking buffer or with blocking buffer without HK2. HK2 binding was detected with anti-Myc-HRP conjugated antibody, and an HRP-catalyzed reaction with a chromogenic substrate solution. Right panel: Nickel-coated plates were incubated with His- PRKAR1a (50 nM) overnight and then with Myc-HK2 (32 nM) for 2h. Thirty minutes before the end of the later incubation, increasing concentrations of G6P (0 – 100uM) were added to the wells and detection was carried out as above. **b.** Glutathione-coated 96-well plates were incubated overnight at 4°C with GST-GSK3b (50 nM) and then incubated with Myc-HK2 (0.25– 64 nM) in blocking buffer or with blocking buffer without HK2. HK2 binding was detected as described above. Right panel: Glutathione -coated plates were incubated with GST-GSK3b (50 nM) overnight and then with Myc-HK2 (32 nM) for 2h. Thirty minutes before the end of the later incubation, increasing concentrations of G6P (0 – 100uM) were added to the wells and detection was carried out as above. **c.** Nickel-coated 96-well plates were incubated overnight at 4°C with His-PRKAR1a (50 nM) or with blocking buffer in absence of PRKAR1a, and then incubated with Myc-HK2 (32 nM) in blocking buffer or with blocking buffer without HK2 for 2h. GST-GSK3b or blocking buffer without GSK3b was then added for 2h. GSK3b binding was detected with anti-GST-HRP conjugated antibody, and an HRP-catalyzed reaction with a chromogenic substrate solution. Right panel: Nickel -coated plates were successively incubated with His-PRKAR1a (50 nM) overnight, HK2-Myc (32nM) for 2h and then GST-GSK3b (32 nM) for 2h. Thirty minutes before the end of the later incubation, increasing concentrations of G6P (0 – 1000uM) were added to the wells and detection was carried out as above. Results are the mean ± SEM of 3 independent experiments. **e.** Nickel-coated plates were incubated overnight at 4°C with His-HK2 (50 nM) and then successively incubated with Myc-PRKAR1α (32nM), PKAc (20nM) and GST-GSK3β (32nM). For the negative control experiments, His-HK2 or Myc-PRKAR1α were omitted. Once all proteins are bound, antibodies against HK2, PRKAR1α, PKAc or GSK3β are added for overnight incubation and detection is carried out after incubation with HRP-conjugated anti-rabbit or mouse IgG as described in Methods. **f.** Nickel-coated plates were incubated overnight at 4°C with His-HK2 (50 nM) and then successively incubated with Myc-PRKAR1α (32nM), PKAc (20nM) and GST-GSK3β (32nM). Kinase buffer in presence or absence of cAMP is added to appropriate wells and P-GSK3β antibody is added overnight, and then detected by incubation with HRP-conjugated anti-rabbit IgG. Bound total GSK3β is detected with anti-GST-HRP conjugated antibody. Both detections were revealed by HRP-catalyzed reaction with a chromogenic substrate solution. For G6P inhibition, G6P was added 30 minutes before the end of the incubation with GST-GSK3β.

The majority of AKAPs have dual specificity for the regulatory subunits of PKA, RIa and R2a, whereas a subset of AKAPs bind RIa only ^27^. We subjected the complex of HK2 and RIa to the disruptor FMP-API-1, which disrupt either R1a or R2a from the dual specificity AKAPs ^28^. We found that FMP-API-1 could not disrupt HK2-RIa interaction in vitro (Extended Data Fig. 7), suggesting that HK2 is a RIa specific AKAP.

### Glucose flux could determine the effect of HK2 on GSK3 phosphorylation

The results described above showed that in the presence of glucose, HK2 elevates GSK3 phosphorylation through its interaction with GSK3 and PKA (which phosphorylates GSK3). This effect on GSK3 phosphorylation is independent of either hexokinase activity or binding to the mitochondria. However, the replacement of glucose with 2-DG inhibited GSK3 phosphorylation through a mechanism that was dependent on hexokinase activity and the phosphorylation of 2- DG to 2-DG6P. Unlike G6P, 2-DG6P is not utilized in glycolysis and therefore accumulates and binds HK2 to elicit conformational changes that promote the dissociation of GSK3 and RIa. If this hypothesis is correct, it is expected that a reduction in glucose metabolism flux in a way that causes G6P accumulation should reduce GSK3 phosphorylation. Therefore, we inhibited the flux of glucose metabolism by exposing the cells to 6-aminonicotinamide (6-AN). 6-AN inhibits 6- phosphogluconate dehydrogenase (6-PGDH), which results in the accumulation of 6-phosphogluconate (6-PG). 6-PG is a competitive inhibitor of phosphoglucose isomerase (PGI), and its inhibition is known to induce G6P accumulation ^29, 30, 31^ (Fig. 4a). Indeed, we found a marked reduction in GSK3*β*phosphorylation following treatment with 6-AN (Fig. 4b, e, and Extended Data Fig. 13b). Interestingly, dehydroepiandrosterone (DHEA), which inhibits glucose- 6-phosphate dehydrogenase (G6PDH) and the first step of the PPP (Fig. 4a), did not inhibit GSK3*β* phosphorylation (Fig. 4b), suggesting that G6P does not sufficiently accumulate if only the first step of the PPP is inhibited. Consistently, we found accumulation of G6P in the cells only after 6- AN treatment and not after DHEA treatment (Fig. 4c). To further corroborate these pharmacological results, we used A549 cells expressing doxycycline (DOX)-induced shRNA targeting G6PDH, PGI or 6PGDH. First, we found that both the pentose phosphate pathway (PPP) and glycolysis were inhibited by either 6-AN or 6PGDH knockdown (Fig. 4d), consistent with inhibition of PGI and the accumulation of G6P. Second, and as expected PGI knockdown decreased secreted lactate, but the secreted lactate was also decreased by 6PGDH knockdown and not by G6PD knockdown further supporting the notion that 6PGDH deficiency via 6-PG accumulation inhibits PGI (Extended Data Fig. 8). Consistent with the pharmacological results, only the knockdown of 6PGDH inhibited the phosphorylation of GSK3*β*, similar to 6AN treatment (Fig. 4e). Furthermore, the same as with 6-AN, 6PGDH deficiency and not PGI deficiency induced the accumulation of both 6PG (Fig. 4f) and G6P (Fig. 4g).

**Figure 4:**
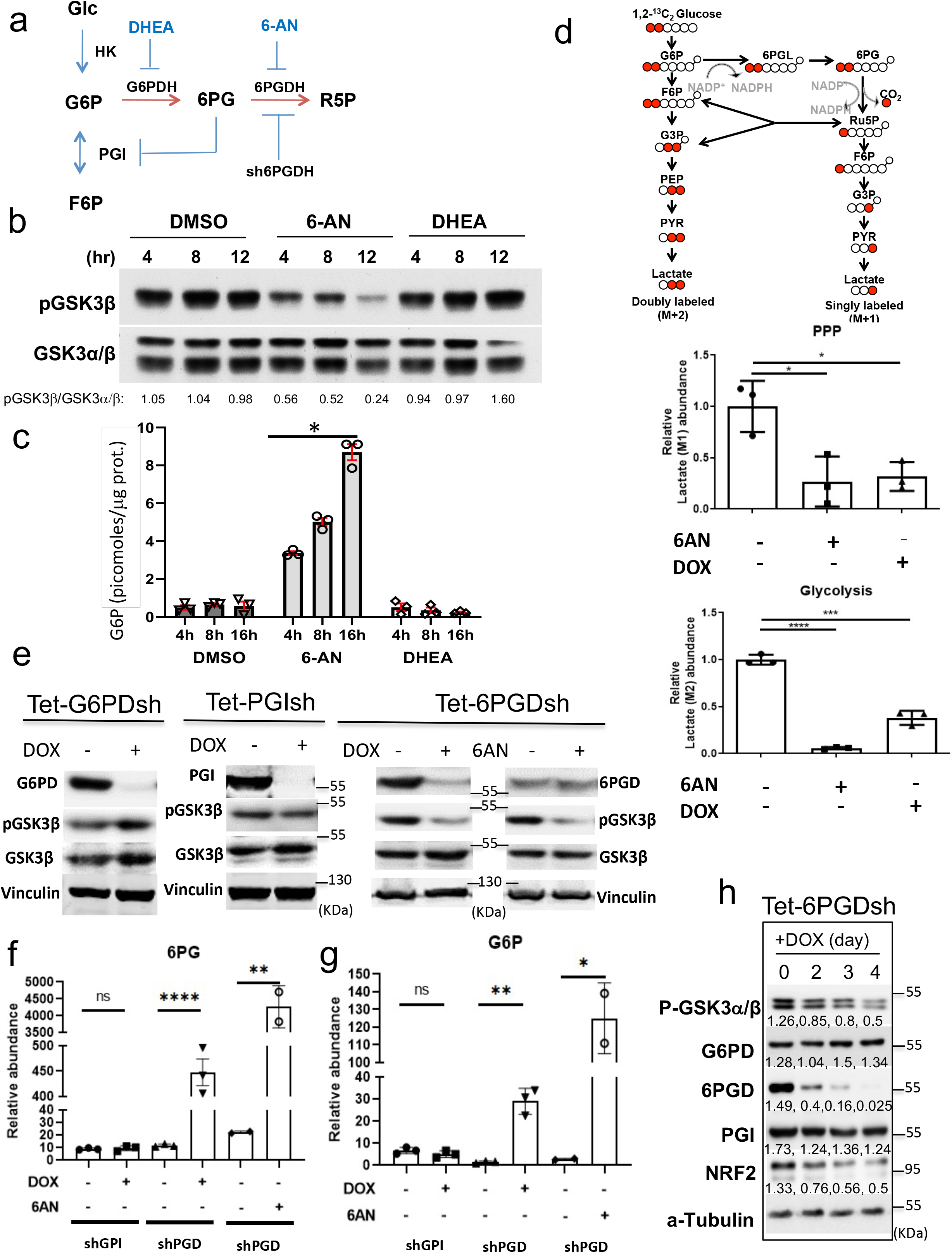
Evidence that intracellular G6P accumulation inhibits GSK3 phosphorylation. **a.** Schematic showing the effect of DHEA, 6-AN, and the knockdown of 6PGDH on the PPP and glycolysis. **b.** Hela cells were treated with either DMSO, 6-AN or DHEA. At the indicated time points, cells were harvested for immunoblotting using anti-p-GSK3β and anti-GSK3*α/β*. **c.** Intracellular levels of G6P in control (DMSO treated), 200μM 6-AN treated, and 200μM DHEA treated cells at different time points after treatment. Results are the mean ± SEM of 3 independent experiments in triplicate. *p<0.05 **d.** Upper panel: Simplified schematic of steps in glycolysis and the pentose phosphate pathway (PPP), showing ^13^C labelling patterns resulting from [1,2-^13^C_2_]-glucose substrate and the conversion to M+1 and M+2 lactate. Red filled circles indicate ^13^C atoms. Abbreviations: G6P, glucose-6-phosphate; 6PGL, 6-phosphogluconolactone; 6PG, 6-phosphogluconate; Ru5P, ribulose-5-phosphate; F6P, fructose-6-phosphate; G3P, glyceraldehyde-3-phosphate; PEP, phosphoenolpyruvate; PYR, pyruvate. Bottom panels: [1, 2-^13^C_2_]-glucose metabolic labelling in A549 cells showing the effect of 6-AN or the DOX-inducible knockdown of 6PDGH on the PPP and glycolysis (cells were treated with either 6-AN for 16h or with DOX for 4 days). Results show relative abundance of intracellular 1x^13^C lactate (M+1) and 2x^13^C lactate (M+2) after 25 mM [1, 2-^13^C_2_]-glucose labeling for 4 h. Results are the mean ± SEM of 3 independent experiments. *p<0.05, ***p<0.01. **e.** A549 cells stably expressing each shRNA in the tet-on system were treated with doxycycline (DOX, 0.2ug/ml) for six days, cells were harvested and analyzed by immunoblotting. A549 cells expressing inducible 6PGD shRNA (Tet-6PGDsh) were also treated with 6-AN for 16h in the absence of DOX. **f.** Cells were treated and isotopically labelled as in d, and 6PG level was measured after GPI and 6PGD knockdown or after treatment with 6-AN. Relative abundance of intracellular 2x^13^C G6P (M+2) and 2x^13^C 6PG (M+2) after 25 mM [1, 2-^13^C_2_]-glucose labeling for 4 h is shown. Results are the mean ± SEM of 3 independent experiments **g.** Cells were treated and isotopically labelled as in d, and G6P level was measured after GPI and 6PGD knockdown or after treatment with 6-AN. **h.** A549 cells expressing inducible 6PGD sh RNA (Tet-6PGDsh) were exposed to DOX and pGSK3 and NRF2 levels were followed for 4 days after DOX addition (Numbers indicate level relative to tubulin).

Since NRF2 expression is elevated in A549 cells because of a lack of functional KEAP1, NRF2 protein stability is controlled in these cells by GSK3 phosphorylation in a Keap1- independent manner ^32^. Consistently, we found that the knockdown of HK2 in A549 cells decreased the phosphorylation of GSK3*α* and GSK3*β* with concomitant decrease in MCL1 and NRF2 protein levels (Extended Data Fig. 9). Thus, it is expected that NRF2 protein levels would be decreased upon depletion of 6-PGDH. Indeed, we found that following the knockdown of 6- PGDH, NRF2 protein levels declined (Fig. 4h).

Collectively, these results show that HK2 acts as a scaffold that brings together PKA and its substrate, GSK3, to facilitate the phosphorylation of GSK3. As depicted in Extended Data Fig. 10, upon accumulation of G6P or 2-DG6P, which induces an allosteric conformational change in HK2, GSK3 is released and subjected to dephosphorylation by PP2A. We concluded that HK2 inhibits GSK3 activity by sequestration and by facilitating its phosphorylation by PKA as was shown for other two AKAPs. Both AKAP200 and GSK3b interacting protein (GSKIP) were shown to interact with both GSK3b and PKA to facilitate the phosphorylation of GSK3b by PKA ^26, 33, 34^ (see Discussion).

### Systemic HK2 deletion in a highly metastatic mouse model of breast cancer decreased metastasis to the lung

The induction of HK2 expression in cancer cells distinguishes them from normal cells. We previously showed that systemic deletion of HK2 in adult mice impeded lung cancer without adverse consequences, suggesting that HK2 could be a selective therapeutic target for cancer ^35^. HK2 was also implicated in breast cancer metastasis ^36^; therefore, we examined whether systemic HK2 deletion could inhibit breast cancer metastasis.

As shown previously ^35^, and in Fig. 6d, HK2 expression was markedly induced in the MMTV-PyMT mouse model of breast cancer metastasis ^37^, thus making it an ideal mouse model to study HK2’s role in metastasis. Therefore, we crossed the *MMTV-PyMT* mice with *HK2^f/f^;UBC- CRE^ERT^*^2^ mice to enable systemic deletion of HK2 after primary breast tumor onset in order to emulate drug therapy and test the effect of systemic HK2 deletion on breast cancer metastasis (Fig. 5a). Once a primary breast tumor was detected by palpation in experimental *MMTV-PyMT; HK2^f/f^; UBC-Cre^ERT^*^2^ mice and control *MMTV-PyMT;HK2^f/f^* mice, the mice were injected with tamoxifen for 7 consecutive days to systemically delete HK2 in the experimental *MMTV-PyMT; HK2^f/f^; UBC-Cre^ERT^*^2^ mice (Fig. 5a). Systemic HK2 deletion increased the time required to reach end-point (when a tumor reached ∼2cm in diameter) (Fig. 5b). At the tumor end-point, the mice were euthanized, and the lungs were analyzed for metastases. As shown in Fig. 5c, lung metastatic lesions were markedly decreased by the systemic deletion of HK2. To determine whether the effect of HK2 on metastasis is cell autonomous, mammary epithelial cells were isolated from the breast tumors of donor *MMTV-PyMT; HK2*^f/f^ mice and established in tissue culture. The cells were infected with GFP-Cre adenovirus to delete HK2, or GFP adenovirus as a control. The cells were transplanted into the mammary fat pad of recipient NOG mice and followed until end-point. HK2 deletion significantly decreased metastatic lung lesions (Fig. 5d).

**Figure 5:**
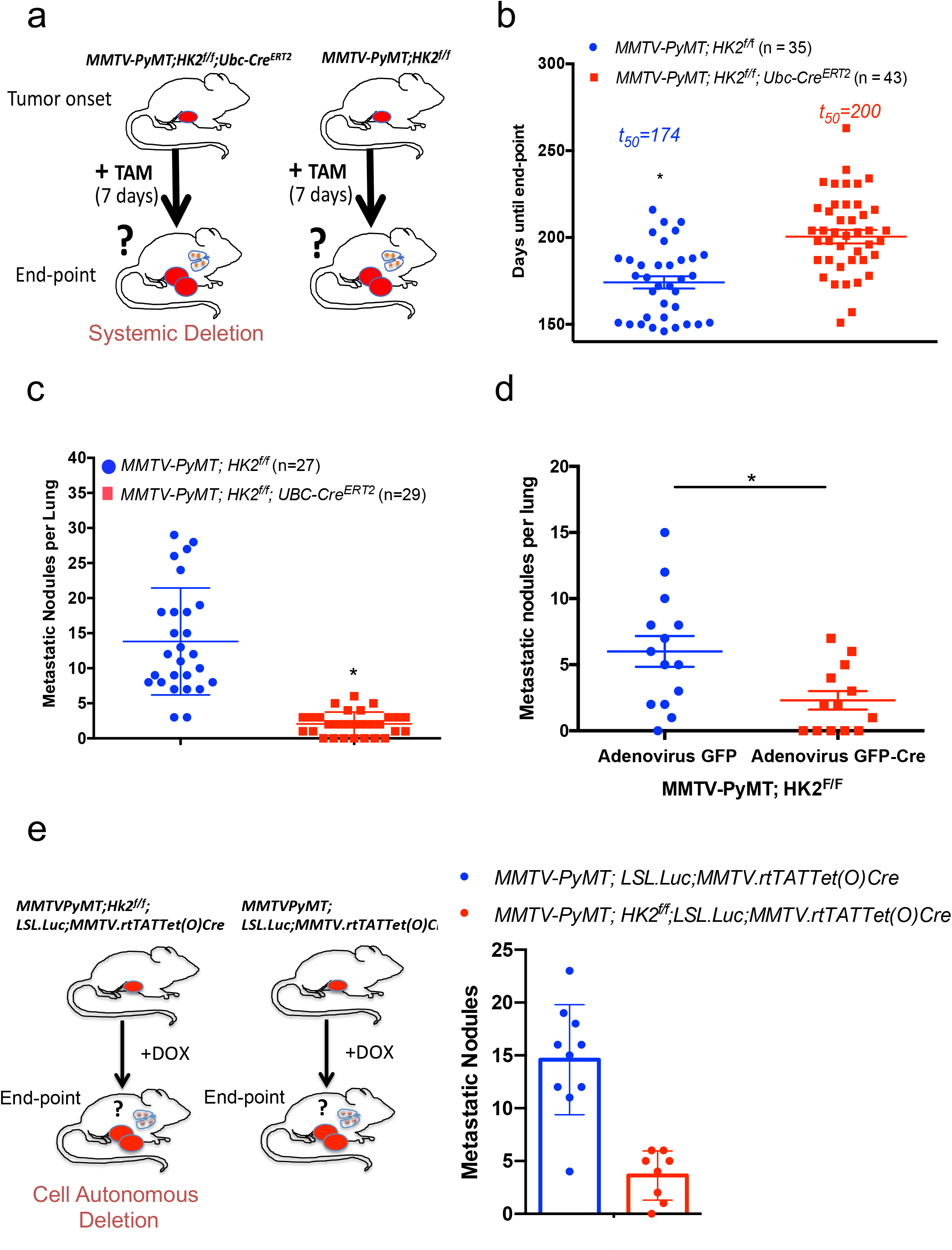
Systemic and cell autonomous deletion of HK2 inhibits metastasis to the lung in *MMTV-PyMT* mice. **a.** Schematic illustration depicting the experimental approach for systemic deletion of HK2 in *MMTV-PyMT* mouse model. Once a primary tumor was palpable, tamoxifen (TAM) was injected for 7 consecutive days in *MMTV-PyMT; HK2^f/f^; UBC-Cre^ERT2^* and *MMTV-PyMT;HK2^f/f^* mice. At endpoint mice were euthanized and lung metastasis was quantified. **b.** Tumor end point analysis in *MMTV-PyMT; HK2^f/f^; UBC-Cre^ERT2^* and *MMTV-PyMT; HK2^f/f^* mice with TAM injection at tumor onset. Results are the mean ± SEM. (p < 0.0001). **c.** Incidence of metastatic lesions in the lungs. When tumors reached end-point, the mice were euthanized, and metastatic lesions in the lungs were quantified. Results are the mean ± SEM. *p < 0.001. **d.** Cell-autonomous deletion of HK2 impairs lung metastasis. Isolated *MMTV-PyMT;HK^f/f^* mammary tumor cells were treated with GFP-Cre adenovirus to delete HK2 or GFP adenovirus as a control. After 72 hr, the cells were transplanted into the mammary fat pads of recipient immunodeficient NOG mice. At the end point, mice were euthanized, and lung metastatic lesions were quantified. The results are the mean ± SEM, *p = 0.0013. **e.** Left: Schematic showing experimental design: *MMTV- PyMT;HK2^f/f^;LSL.Luc;MMTV.rtTATet(O)Cre* and control *MMTV- PyMT;LSL.Luc;MMTV.rtTATet(O)Cre* mice were subjected to DOX diet after tumor onset and metastasis was quantified at end point. Right: Quantification of metastasis to the lung. The results are the mean ± SEM. **p=0.0038.

**Figure 6:**
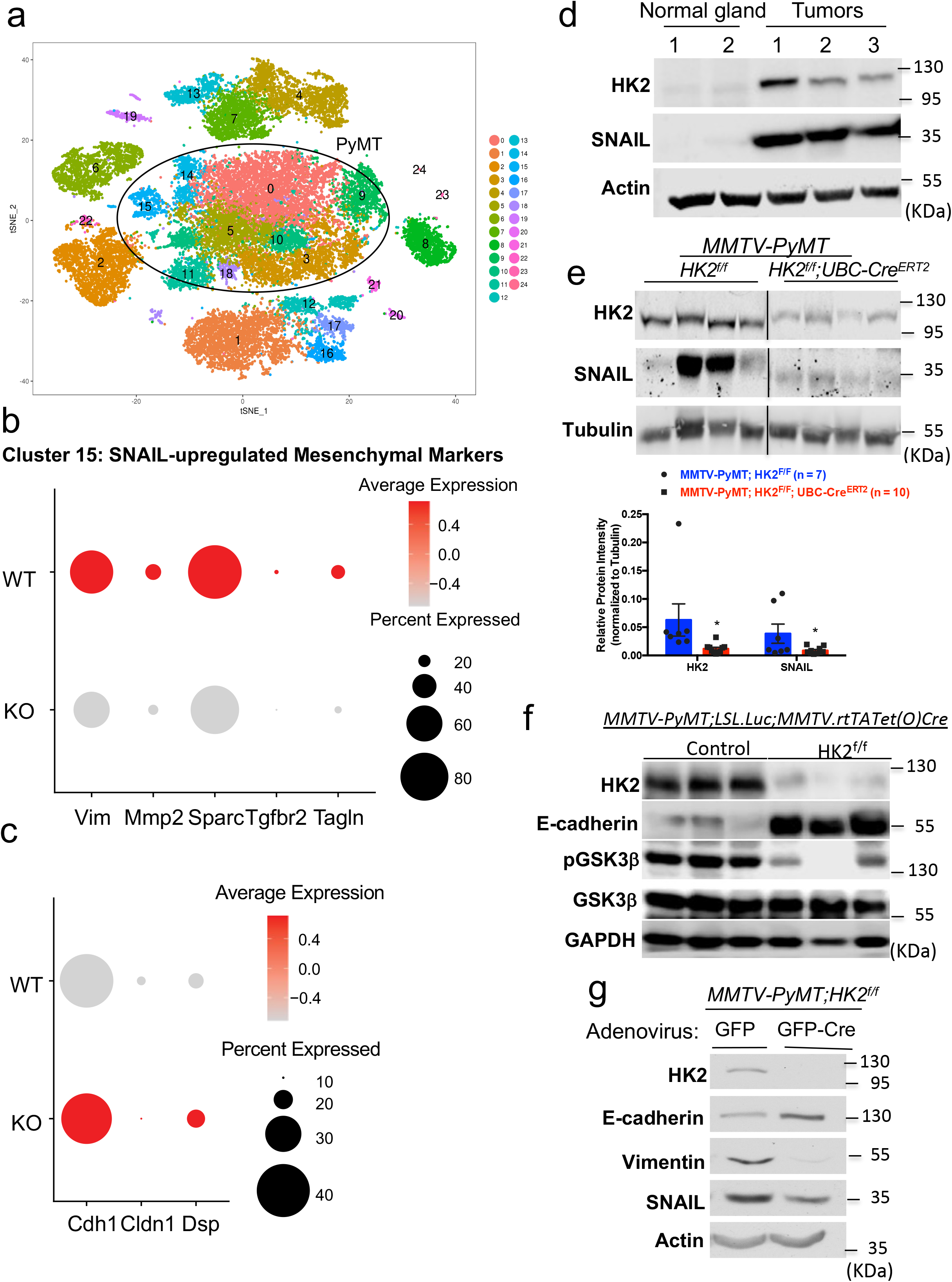
The effect of HK2 deletion on specific SNAIL target gene expression in epithelial and mesenchymal cell clusters and SNAIL protein levels in the primary tumors. **a.** Single cell RNA sequencing and clustering of primary tumor cells derived from tumors at end- point of *MMTV-PyMT;HK2^f/f^* (n=7) and *MMTV-PyMT;HK2^f/f^;UBC-Cre^ERT2^* (n=8) mice after TAM exposure. t-Distributed Stochastic Neighbor Embedding (tSNE) is shown. The color-coded clusters are grouped together by similar gene expression. Twenty-five clusters including nine primary tumor PyMT-expressing clusters were identified. **b.** Dot plot showing the expression of SNAIL targets important for EMT and metastasis (Vim, Mmp2, Sparc, Tfgbr2, Tagln) in Cluster 15 from WT and KO (*MMTV-PyMT;HK2^f/f^* and *MMTV- PyMT;HK2^f/f^;UBC-Cre^ERT2^*) tumors after TAM exposure at tumor onset). High RNA expression is indicated by a dark red color. **c.** Dot plot showing the expression epithelial markers such as E-cadherin (Cdh1), Claudin-1 (Cldn1), and Desmoplakin (DSP) in Cluster 14 from WT and KO (*MMTV-PyMT;HK2^f/f^* and *MMTV- PyMT;HK2^f/f^;UBC-Cre^ERT2^* tumors after TAM exposure at tumor onset . A high RNA expression is indicated by a dark red color. **d.** Immunoblot image of HK2 and SNAIL protein levels in tissue samples collected from normal mammary glands and primary mammary gland tumors. **e.** Upper: Immunoblot image of HK2 and SNAIL protein levels in tissue samples collected from primary mammary gland tumors at end-point after exposure to TAM at tumor onset, in *MMTV- PyMT;HK2^f/f^* and *MMTV-PyMT;HK2^f/f^;UBC-Cre^ERT2^* mice. Bottom: Quantification of HK2 and SNAIL protein levels, *MMTV-PyMT;HK2^f/f^* (n=7), *MMTV-PyMT;HK2^f/f^;UBC-Cre^ERT2^* (n=10). Results are the mean ± SEM. *p≤0.05. **f.** Immunoblot image of HK2, pGSK3b, GSK3b, E-cadherin, and SNAIL protein levels in primary mammary tumors at end-point derived from *MMTV-PyMT;LSL.Luc;MMTV.rtTATet(O)Cre* control mice (n=3) or *MMTV-PyMT;HK2^f/f^;LSL.Luc;MMTV.rtTATet(O)Cre* mice (n=3)subjected to DOX diet after tumor onset. **g.** Representative immunoblot image of HK2, E-cadherin, vimentin, and SNAIL protein levels in protein extracts from cells isolated from tumors in *MMTV-PyMT;HK2^f/f^* mice after infection with adenovirus expressing either GFP or GFP-Cre (n*≥*3).

To further determine the cell autonomous effect of HK2 on metastasis, we generated *MMTV-PyMT;Hk2^f/f^;LSL.Luc;MMTV.rtTATet(O)Cre* mice in which HK2 could be specifically deleted in the mammary gland immediately after tumor onset by exposing the mice to doxcycyline diet (Fig. 5e), and found a marked reduction in the metastatic lesions in the lungs. Thus, the effect of HK2 on metastasis is, at least in part, cell autonomous.

### HK2 deletion decreased the expression of epithelial mesenchymal transition (EMT) genes and SNAIL protein abundance

We analyzed the primary tumors with single-cell RNA sequencing (scRNA-seq). We adopted Drop-seq technology for scRNA-seq as previously described ^38, 39–41^. We sequenced 28417 cells from 15 biological replicates of primary tumors derived from *MMTV-PyMT;HK2^f/f^;UBC- Cre^ERT^*^2^ mice with and without systemic HK2 deletion (7 replicates and 8 replicates, respectively). We found 25 clusters and identified 9 clusters (0, 3, 5, 9, 10, 11, 14, 15, and 18) within the primary tumor based on the expression of PyMT (Fig. 6A, and Supp. Table 1). However, the percentage of cells from the primary tumors after the systemic deletion of HK2 that grouped into the 9 PyMT clusters was not significantly different from that in the control primary tumors (Extended Data Fig. 11, and Supp. Table 1). However, when we had a close look at two adjacent clusters; cluster 14, which expresses the highest levels of epithelial markers, and cluster 15, which expresses the highest levels of mesenchymal markers, we found changes in the expression of SNAIL target genes by the deletion of HK2. Cluster 14 exhibited high expression of the epithelial markers including E-cadherin (Cdh1), Claudin-1 (Cldn1) and Desmoplakin (Dsp), which are known to be repressed by SNAIL ^42–44^. Cluster 15 expresses the highest level of genes that promote EMT and are known SNAIL targets (vimentin (Vim), Matrix metaloproteinase 2 (Mmp2), secreted protein acidic and cysteine rich (Sparc), TGF beta receptor 2 (Tgfbr2), and Transgelin (Tagln)). Interestingly, the systemic deletion of HK2 markedly reduced SNAIL target genes in Cluster 15 and increased epithelial genes in Cluster 14 in comparison with their levels in the control samples (Fig. 6b and c). These results suggest that the loss of HK2 in primary tumors impaired the expression of SNAIL-regulated targets important for EMT and metastasis.

EMT is important for breast cancer metastasis, and the deletion of SNAIL in the mammary gland of MMTV-PyMT mice was shown to inhibit metastasis to the lung ^45, 46^. Therefore, the results raised the possibility that the deletion of HK2 could affect metastasis by impairing SNAIL activity to mediate EMT. When we analyzed the primary tumors for SNAIL protein levels, we first found that HK2 was induced, with a concomitant marked increase in SNAIL in the tumors when compared to that in normal mammary glands (Fig. 6d); second, SNAIL protein levels in the tumors were markedly reduced after HK2 deletion (Fig. 6e). To determine if the effect of systemic HK2 deletion on SNAIL is cell autonomous, we analyzed primary tumors in the cell autonomous mouse model described in Fig. 5e. As shown in Fig. 6f, cell autonomous HK2 deletion in the mammary gland decreased p-GSK3b, with concomitant increase in E-cadherin, in the primary tumors. Finally, when cells were isolated from the primary tumors of *MMTV-PyMT; HK2^f/f^* mice and HK2 was deleted by adenovirus expressing Cre, we found that the SNAIL protein level was decreased, with a concomitant increase in the E-cadherin protein level and a decrease in the vimentin protein level (Fig. 6g). Taken together, these results indicate that HK2 deletion reduced SNAIL protein level, and leading to a decrease in its transcriptional activity important for EMT.

### Increased HK2 expression correlates with increased metastatic potential

To further investigate the cell autonomous role of HK2 in metastasis, we utilized two isogenic mammary tumor cell lines (67NR and 4T1) that, when implanted into the mammary gland of mice, display different metastatic potential ^47^. Both cell lines have the ability to form mammary tumors within a month. However, the 67NR cells only forms primary tumors with no metastasis, whereas the 4T1 cells forms primary tumor with high incidence of metastasis to the lung. When HK2 protein levels were analyzed, the non-metastatic cell line 67NR exhibited markedly lower levels when compared to the highly metastatic 4T1 cell line (Fig. 7a). As a result, this enabled us to manipulate the level of HK2 expression in these cell lines to determine whether HK2 plays a cell autonomous role in the metastatic potential of these cell lines. Therefore, we silenced HK2 in 4T1 cells (4T1shHK2) and overexpressed HK2 in 67NR cells (67NR HK2) (Extended Data Fig. 12a). The knockdown of HK2 in 4T1 cells decreased transwell migration, and invasion (Extended Data Fig. 12b, c) while the overexpression of WT HK2 in 67NR cells increased migration (Extended Data Fig. 12b). Interestingly, even overexpression of kinase inactive HK2DA mutant in 67NR cells was able to increase migration (Extended Data Fig. 12c), suggesting that, at least in these cells, the scaffolding activity of HK2 is pre-dominant in promoting migration. Thus, overall, HK2 level appears to correlate with the metastatic potential of these cell lines. Next, we assessed the effect of HK2 on the metastasis of 4T1 cells in vivo. 4T1 or 4T1 shHK2 cells were injected orthotopically into the fourth mammary fat pad of syngeneic mice. At end-point, when primary tumors reached the same size, the mice were euthanized, and the lungs were analyzed for lung metastases. The knockdown of HK2 significantly decreased the number of lung metastases. The control 4T1 tumors had an average of 28 lung metastases whereas the mice with 4T1 shHK2 tumors had an average of 4 lung metastases (Fig. 7b).

**Figure 7.**
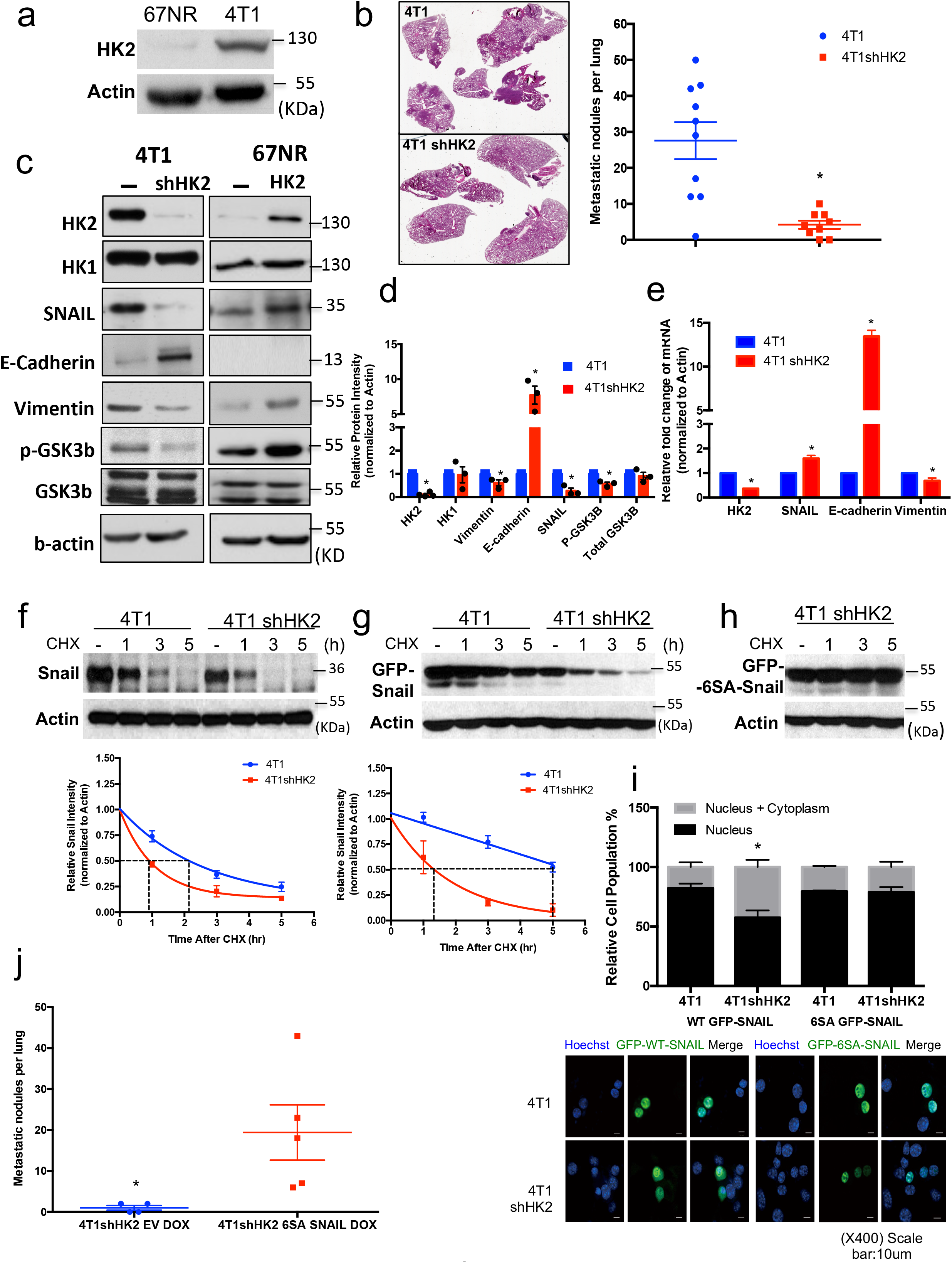
HK2 silencing in 4T1 cells inhibits metastasis to the lung and impairs EMT gene expression, and SNAIL protein stability and nuclear localization in a GSK3-dependent manner. **a.** Immunoblot image showing HK2 levels in two isogenic breast cancer cell lines: 67NR (a non-metastatic line) and 4T1 (a highly metastatic line). **b.** Left: Representative images showing H&E staining of lung sections after orthotopic transplantation of 4T1 or 4T1shHK2 cells into Balb/cj mice. Right: Quantification of lung metastatic lesions (4T1 transplantation, n=10; 4T1 shHK2 transplantation, n=9) at end point. Results are the mean ± SEM. *p<0.001 **c.** A representative immunoblot image showing the effect of HK2 silencing in 4T1 cells and overexpression of HK2 in 67NR cells on SNAIL, E-cadherin, and vimentin protein levels and GSK3*β* phosphorylation (n=3). **d.** Quantification of protein levels after HK2 silencing in 4T1 cells. Results are the mean ± SEM of 3 independent experiments. *p<0.05. **e.** Quantitative RT-PCR analysis measuring relative mRNA levels in 4T1 and 4T1shHK2 cells. Results are the mean ± SEM (n=5). *p< 0.03. **f.** Protein stability of SNAIL in 4T1 and 4T1shHK2 cells as measured after exposure to cycloheximide (CHX). Upper: Representative immunoblot image. Bottom: Plot showing SNAIL protein half-life after quantification relative to b-actin in 3 independent experiments. The half-life results are the mean ± SEM. p<0.01 **g.** Protein stability of transiently expressed GFP tagged SNAIL in 4T1 and 4T1shHK2 cells as measured after exposure to cycloheximide (CHX). Upper: Representative immunoblot image. Bottom: Plot showing GFP-SNAIL protein half-life after quantification relative to b-actin in 3 independent experiments. The half-life results are the mean ± SEM. p<0.05 **h.** Immunoblot image showing the level of transiently expressed GFP-tagged 6SA-SNAIL mutant in 4T1shHK2 after exposure to cycloheximide over a 5-hr time course. **i.** GFP-SNAIL and GFP-6SA-SNAIL were transiently expressed in 4T1 and 4T1shHK2 cells. After fixation of the cells, the localization of SNAIL (green) and nuclei (blue; Hoechst 33342) was examined under a fluorescence microscope and quantified (bottom). The bar graph represents the relative GFP-SNAIL localization in the cytoplasm versus in the nucleus in 4T1 and 4T1shHK2 cells. Results are the mean ± SEM, n=3. *p< 0.05. **j.** Transient expression of 6SA SNAIL in 4T1shHK2 cells restores metastasis to the lung. 4T1shHK2 cells expressing DOX-inducible empty vector (EV) and 4T1shHK2 cells expressing DOX-inducible 6SA SNAIL were orthotopically implanted in Balb/cj mice (4T1shHK2 EV, n=4 and 4T1shHK2 10 6SA SNAIL n=5), and exposed to DOX for one week. At end point lung metastatic lesions were quantified. The results are the mean ± SEM. *p<0.05

### HK2 regulates the protein stability of the EMT transcription factor SNAIL in a GSK3β-dependent manner

Based on the results in Fig. 6, we concluded that HK2 deletion might affect EMT and metastasis by downregulating SNAIL protein levels. Because SNAIL protein level was shown to be regulated by GSK3 phosphorylation, targeting it for proteasomal degradation ^18^, we determined whether HK2 affects the level of SNAIL via GSK3.

Consistent with the results obtained with the primary MMTV-PyMT tumors (Figs. 6e- g), SNAIL protein levels were decreased by knocking down HK2 in 4T1 cells, with a concomitant decrease in Vimentin and increase in E-cadherin at both the protein and mRNA levels (Figs. 7c-e). Interestingly, the mRNA level of SNAIL (Fig. 7e) did not correlate with SNAIL protein levels (Fig. 7c), indicating that SNAIL protein stability was impaired by HK2 silencing. We also found that the decrease of SNAIL protein level in 4T1shHK2 cells was associated with a concomitant decrease in GSK3β phosphorylation, whereas the increase in SNAIL protein level in 67NR WTHK2 cells is associated with concomitant increase in GSK3β phosphorylation (Fig. 7c). To directly test the possibility that HK2 affects SNAIL protein stability, we analyzed the half-life of SNAIL in 4T1 cells after the knockdown of HK2 and found that the half-life of SNAIL protein was markedly decreased (Fig. 7f). The half-life of ectopically expressed SNAIL-GFP fusion protein was also markedly decreased by the knockdown of HK2 (Fig. 7g), but this was not true for the half-life of 6SA-SNAIL-GFP fusion protein, in which all the serine residues that are phosphorylated by GSK3 were converted alanine residues ^18^ (Fig. 7h).

GSK3β not only targets SNAIL for degradation, but also shifts the localization of SNAIL from the nucleus to the cytoplasm ^18^. We wanted to determine whether HK2 expression could mediate the localization of SNAIL in a GSK3β-dependent manner. We therefore quantified the intracellular localization of WT SNAIL-GFP fusion protein or mutant 6SA- SNAIL-GFP fusion protein, which is resistant to GSK3 phosphorylation ^18^, in 4T1 and 4T1shHK2 cells. WT SNAIL was mainly localized to the nucleus in the 4T1 cells, but its localization shifted to a more cytoplasmic localization in the 4T1shHK2 cells, whereas the SNAIL-6SA mutant remained mostly nuclear in both cell lines (Fig. 7i). Taken together these results strongly demonstrate that HK2 affects SNAIL protein stability and nuclear localization via its effect on GSK3. Furthermore, 2DG treatment, which led to the accumulation 2DG6P, as well as 6-AN treatment that leads to accumulation of G6P and activation of GSK3 decreased SNAIL protein levels in 4T1 cells (Extended Data Fig. 13 a, b, c). Finally, the same as was shown for HEK293 cells in Fig. 2f, we found that in 4T1 cells endogenous HK2 binds endogenous GSK3 only in the presence of glucose and not in the in the presence of 2-DG (Extended Data Fig. 13c).

If the HK2 effect on metastasis is via SNAIL, it is expected that 6SA-SNAIL mutant would restore metastasis in HK2-deficient cells. Since it was reported that only transient SNAIL expression induces metastasis, whereas constitutive expression of SNAIL does not and may impede metastasis ^46^, we generated a 4T1shHK2 cell line with doxycycline-inducible 6SA-SNAIL expression. Transient inducible expression of SNAIL-6SA mutant in the 4T1shHK2 in mice rescued changes in lung metastasis (Fig. 7j). We concluded that HK2 affects metastasis through its effect on SNAIL protein stability and nuclear localization.

### HK2 deficiency impaired global O-GlcNAcylation and specific SNAIL O-GlcNAc modification

It was previously reported that O-linked-N-acetylglucosamine (O-GlcNAc) modification (O-GlcNAcylation) is important to breast cancer primary tumor progression and metastasis, and that primary tumor GlcNAcylation levels were found to be increased during mammary tumor progression in MMTV-PyMT mice ^48^. Uridine 5’-diphospho-N acetylglucosamine (UDP-GlcNAc), which is transferred to the serine or threonine residues of proteins to yield O-GlcNAcylation, is generated by the hexosamine biosynthetic pathway. Flux in the hexosamine biosynthesis pathway is dependent on G6P, raising the possibility that HK2 deficiency reduces glycosylation. By using an O-GlcNAc specific antibody on total protein lysates, we found that the systemic deletion of HK2 in MMTV-PyMT mice decreased the total O-GlcNAcylation levels in the primary tumors at end-point (Fig. 8a). Consistently, the nonmetastatic 67NR cells, which express relatively low level of HK2, had less total O-GlcNAcylation when compared to the metastatic 4T1 cells, which express higher level of HK2 (Fig. 8b). Furthermore, the knockdown of HK2 in 4T1 cells decreased protein O-GlcNAcylation, and overexpression of WT HK2 in 67NR cells increased the O-GlcNAcylation (Fig. 8b). Taken together these results strongly suggest that HK2 deficiency reduces global O-GlcNAcylation. To further interrogate the role of HK2 in the hexosamine pathway we conducted a metabolic labeling experiment to determine whether HK2 deficiency decreases the flux of glucose to the hexosamine pathway. As shown in Fig. 8C, HK2 deficiency markedly decreased the incorporation of U^13^ C_6_ glucose in all UDP-GlcNAc isotopomers.

**Figure 8.**
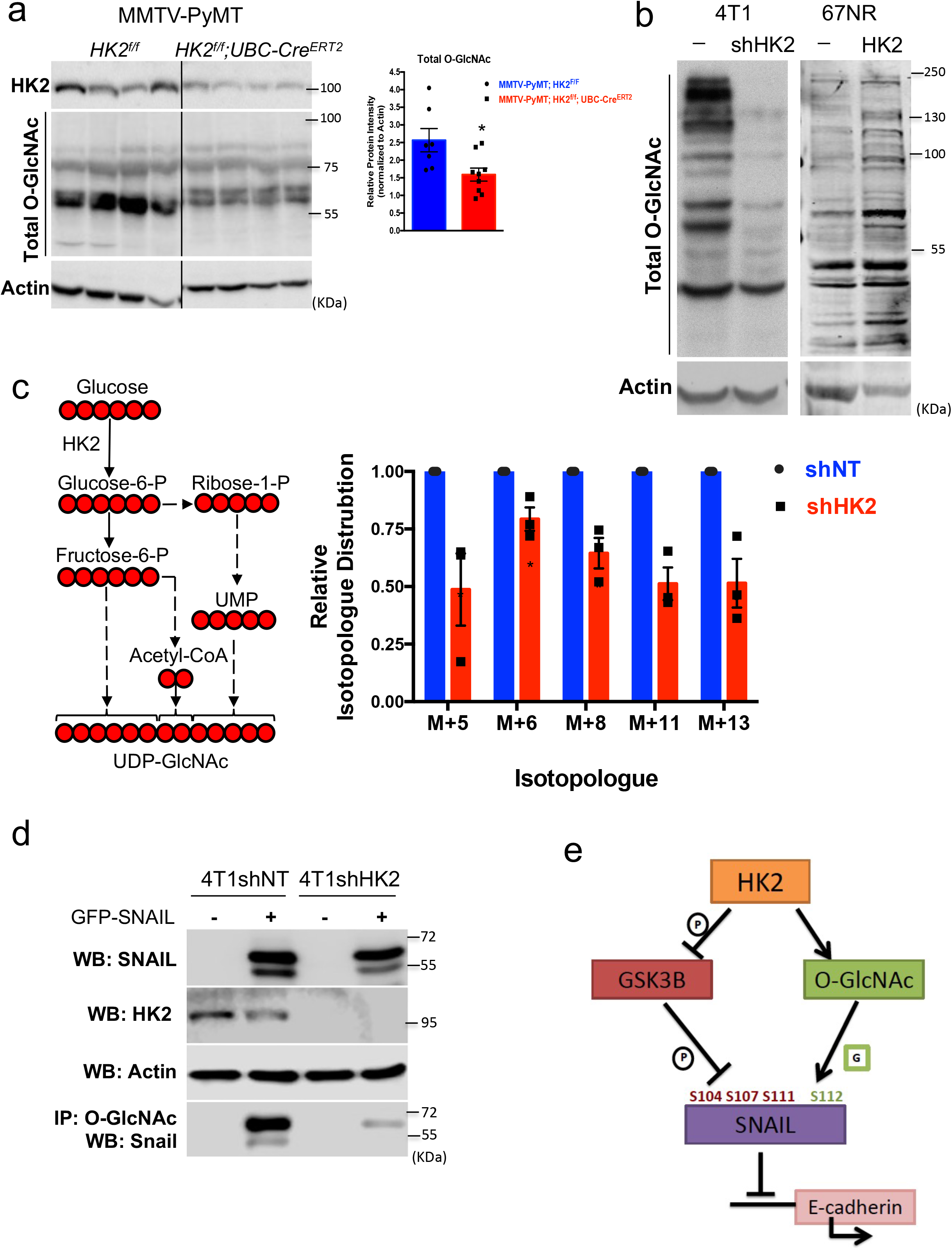
HK2 deficiency inhibits the incorporation of metabolically labeled glucose into UDP-N-acetylglucosamine and reduces total and SNAIL O-GlcNAc modification. **a.** Left: Immunoblot images showing total O-GlcNAc protein modification in MMTV-PyMT mammary gland tumors after systemic deletion of HK2 (n=4). Right: Quantification of total O-GlcNAc modifications in control primary tumors and primary tumors after systemic HK2 deletion (n=9). Results are the mean ± SEM. *p< 0.05. **b.** Immunoblot images of total O-GlcNAc protein modification in 4T1 cells after HK2 silencing and 67NR cells after HK2 overexpression. **c.** Tracing of UDP-GlcNAc isotopomers after culturing 4T1 and 4T1shHK2 cells with 5mM of U^13^C_6_-glucose for 5min. Tracing was performed by ion chromatography (IC-MS). Data shown as fold changes relative to the control 4T1shNT cells for each isotopomer for UDP-GlcNAc with natural abundance correction and represented as means SEMs *p < 0.05 from triplicate experiments using an unpaired t-test (n=3). The schematic shows the contribution U^13^C_6_- glucose-6-P to the UDP-GlcNAc isotopomers. **d.** Immunoprecipitation with anti-O-GlcNAc and immunoblotting with anti-SNAIL after transient transfection of 4T1 and 4T1shHK2 cells with GFP-SNAIL. **e.** Schematic depicting how HK2 affects SNAIL protein stability and activity.

O-GlcNAcylation had been shown to contribute to SNAIL protein stability by blocking GSK3β from serially phosphorylating SNAIL to promote its degradation ^49^. O-GlcNAc at Ser112 stabilizes SNAIL by blocking GSK3β from serially phosphorylating at serine 104, 107, and 111 and promoting SNAIL degradation ^49^. Therefore, we wanted to examine whether in addition to the noncatalytic effect of HK2 on GSK3 activity, HK2 could affect the glycosylation of SNAIL. To verify whether there are specific changes in O-GlcNAcylation of the SNAIL protein itself occurred, GFP- SNAIL was transiently expressed in the 4T1 and 4T1shHK2 cells. This was followed by immunoprecipitation with anti-O-GlcNAc antibody and immunoblotting with anti-SNAIL antibody (Fig. 8d). The results clearly showed that SNAIL O-GlcNAcylation was diminished by the deficiency of HK2. We concluded that in addition to its noncatalytic activity on GSK3, HK2 catalytic ability can also contribute to the stability and nuclear localization of SNAIL via O-GlcNAcylation (Fig. 8e).

## Discussion

HK2 is markedly induced in cancer cells when compared to normal cells. An increase in HK2 expression and activity is a critical determinant of the accelerated glucose metabolism in cancer cells. We previously documented that HK2 is required for tumorigenesis both in vitro and in vivo ^4, 35, 50^. HK2 expression also correlates with the incidence of breast cancer metastasis to the brain^36^. However, thus far the pro-tumorigenic role of HK2 has been attributed to its metabolic activity as a glucose kinase that converts glucose to glucose 6-phosphate. Here we uncovered a new activity of HK2 independent of its catalytic activity. We showed that HK2 binds to GSK3 as well as the PKA regulatory subunit RIa, and it is conceivable that HK2 thereby facilitates the phosphorylation of GSK3 by PKA. Since these interactions may not require HK2 catalytic activity, HK2 may act as purely a scaffold in this scenario. On the other hand, the conversion of glucose to G6P or 2-DG to 2-DG6P by catalytically active hexokinase seems to be required for the diminished phosphorylation of GSK3 and therefore its activation. Binding of G6P or 2-DG6P to HK2 induces a conformational change that can dissociate GSK3. Indeed, we found that 2-DG disrupted the HK2-GSK3 and HK2-RIa interactions. In addition, we found that following the dissociation of GSK3 by 2-DG, it interacts with PP2A, which facilitates its dephosphorylation and thus its activation.

Our results suggest that HK2 can function as an A-kinase-anchoring protein (AKAP), a novel moonlighting function for this important glycolytic enzyme that has not previously been described. Two of the AKAPs, AKAP220 and GSKIP, were shown to facilitate GSK3 phosphorylation by anchoring PKA through its interaction with PRKAR2 ^26, 33, 34^, but AKAP220 was also reported to interact with PRKAR1 ^34^. Interestingly, some AKAPs were shown to interact with phosphatases. For instance, AKAP220 was shown to interact with PP1, which could contribute to the regulation of GSK3 phosphorylation ^34^. Although we were unable to demonstrate binding of PP2A to HK2, we showed that GSK3 binds PP2A when not sequestered by HK2. That said, we cannot completely exclude PP2A binding to HK2. It is interesting to note that D-AKAP1 interacts with the outer mitochondrial membrane through an hydrophobic motif similar to those of HK1 and HK2 ^51^ and thus could be in close proximity to the mitochondrial HK2. Therefore, we could not completely rule out the possibility that PKA and phosphatases associated with D-AKAP1 could also influence the phosphorylation of GSK3 associated with HK2, although we found that a mitochondrial binding-deficient mutant of HK2 could still exert a similar effect on GSK3. We clearly showed that HK2 binds GSK3b and affects GSK3b phosphorylation in a kinase-independent manner. The yeast-two hybrid screen showed that the amino-terminus domain of RIa, which possess the docking site to AKAPs ^24^, is sufficient to bind HK2. The binding of HK2 to RIa was confirmed by co-immunoprecipitation in mammalian cells and together with the inhibition of HK2-mediated GSK3 phosphorylation by pharmacological inhibitors of adenylate cyclase and PKA, the results strongly suggest that HK2 could be a bona-fide AKAP. This is confirmed by the in vitro binding assays with purified proteins that showed unequivocally a direct binding between HK2, GSK3b, and RIa to form complexes that are disrupted upon addition of G6P.

All ∼100 kDa high affinity hexokinases are thought to have arisen from gene duplication and tandem ligation events involving a common ancestral ∼50 kDa hexokinase ^21, 52^. As a consequence of this common evolutionary origin, the individual amino- and carboxy-halves of HK2 are highly homologous with each other and with the corresponding hemidomains of HK1. Given their known structural similarities, it is conceivable that both hemidomains could anchor GSK3, PKA, and possibly PP2A (Extended Data Fig. 10). Although presently speculative, we also cannot exclude the possibility that HK1 may serve similar AKAP-like roles.

In summary, we have discovered a novel phenomenon, wherein glucose metabolites regulate the activity of GSK3 with far-reaching and profound biological significance. HK2 can sequester GSK3 and facilitate its phosphorylation. However, a metabolic slow-down that leads to the accumulation of the most upstream glucose metabolite, G6P, could trigger the dephosphorylation and consequent activation of GSK3. In turn, GSK3 could affect cell proliferation, cell growth, cell survival by multiple mechanisms, and tumorigenesis. We showed that HK2 could affect the levels of three proteins that are known targets of GSK3: MCL1, NRF2, and SNAIL. Here, we also showed for the first time that systemic deletion of HK2 inhibited breast cancer metastasis in a mouse model of breast cancer metastasis. Our results strongly suggest that the effect of HK2 on metastasis is mostly cell autonomous, although we cannot completely exclude non-cell autonomous effects. We have demonstrated that HK2 depletion reduced SNAIL protein levels in a GSK3-dependent manner and in turn affected the expression of EMT genes and metastasis. It was shown that the extent of GlcNAcylation is induced in human breast cancer and is further elevated in metastatic lymph nodes ^53^. It was also reported that O-GlcNAcylation and GlcNAcylation in general are important for breast cancer tumor progression and metastasis ^48, 53^ and that SNAIL phosphorylation by GSK3 could be counteracted by O-GlcNAcylation ^49^.

Therefore, we examined the effect of HK2 depletion on the hexosamine pathway, as well as general and SNAIL-specific O-GlcNAcylation. Our findings showed that HK2 depletion markedly reduced the hexosamine pathway and both general and SNAIL-specific O-GlcNAcylation. Thus, our results showed that both the noncatalytic activity, uncovered here, and the catalytic HK2 activity contribute to the regulation of SNAIL protein levels and metastasis. Importantly, as we had previously suggested the feedback inhibition of HK2 by its own catalytic product G6P could be used as a strategy to target HK2 for cancer therapy by developing mimetics of G6P or 2DG6P to inhibit its activity ^35^. Using this approach, inhibitors that selectively target HK2 and not HK1 were developed ^5^. Since G6P or 2DG6P also dissociates GSK3 from HK2 thereby increasing the activity of GSK3, this approach could inhibit both the glycolytic activity of HK2 as well as its moonlighting activity as a promoter of metastasis.

## Methods

### Cell culture, transfection and transduction

MI5-4 CHO cells were kindly provided by Dr. John Wilson. CHO cells were cultured in α-MEM supplemented with 10% FBS (Gemini) and 2 mM glutamine. All other cells were cultured in DMEM (Invitrogen) containing 10% FBS (Gemini). For glucose starvation, the medium was replaced with glucose-free DMEM (Invitrogen) containing 10% dialyzed FBS (Gemini) after washing with PBS. For immunoprecipitation, 2×10^6^ HEK293-HK2-HA cells were plated in 6-cm dishes one day before transfection using 2 µg of each plasmid. After 24 hr of transfection, cells were used for experiments. For transient transfection of shRNA plasmids in HEK293-HK2-HA cells, 2.5×10^5^ cells were plated in 6-well plates one day before transfection using 3 µg of each shRNA plasmid and Lipofectamine 2000. After 48 hr of transfection, cells were used for experiments. For the tet-inducible shRNA expression system, each shRNA sequence was inserted into the Tet-pLKO-puro vector (a gift from Dmitri Wiederschain; Addgene plasmid #21915). The most efficient shRNA sequences we used were TRCN0000281204 for G6PDsh, TRCN0000274975 for GPGDsh, and TRCN0000290649 for GPIsh. For lentivirus production, each lentiviral vector was cotransfected with pMD2.G and psPAX2 (gifts from Didler Trono; Addgene plasmids #12259 and #12260, respectively) in 293T cells using Lipofectamine 2000 as described in a protocol on the Addgene website. Lentivirus infection was performed overnight in the presence of polybrene, and selection was carried out with 10 µg/ml blasticidin.

The cDNA for rat HK2 was subcloned into the pcDNA3-HA vector. Site-directed mutagenesis of HK2-DA (D209A/D657A; a non-glucose-binding mutant), SA (S155A/S603A; a noncatalytic mutant), and dMT (d1-20aa) was performed using QuickChange II XL (Stratagene) according to the manufacturer’s instructions. The HA-tagged cDNAs were subcloned into the pLenti6-D-TOPO vector (Invitrogen). The cDNA for human GSK3*β*was subcloned into the pCMV- myc vector. cDNA for human PRKAR1a was subcloned into the pLenti6-D-TOPO vector. All lentiviruses were produced using the BLOCK-iT lentiviral expression system (Invitrogen) according to the manufacturer’s instructions.

Polyclonal 4T1 cells with stable HK2 knockdown were generated after infection with pLenti6-puro vector expressing a mouse HK2 hairpin and selection with puromycin. The HK2 mouse target sequence was 5’-GCATATGATCGCCTGCTTAT-3’. Flag-Snail-6SA and GFP-SNAIL-6SA were obtained from Mien-Chie Hung through Addgene (Addgene plasmid # 16225; http://n2t.net/addgene:16225 ; RRID:Addgene_16225) and cloned into the PCW-puro vector (addgene: 50661, following the removal of Cas9 with NHEI and BamHI) using Gibson Assembly. The ViraPower^TM^ lentiviral expression system from Thermo Fischer Scientific was used for virus production. Cells were selected with 2 µg/µl puromycin or 8 µg/µl blasticidin for 5 days. The 4T1 cells were sent for screening with the IDEAX Impact I panel before orthotopic transplantation into mice. The cells used to test for total O-GlcNAc levels, and for immunoprecipitation experiments were plated in 5 mM glucose DMEM with 10% serum for at least 5 days prior to harvesting for experiments.

### Immunoprecipitation and immunoblotting

For immunoprecipitation, cells grown in 6-cm dishes were lysed in IP buffer (50 mM Tris-Cl (pH 8.0), 0.5% NP-40, 150 mM NaCl, 1 mM EGTA, 1 mM EDTA) containing phosphatase inhibitors (10 mM sodium pyrophosphate, 20 mM β-glycerophosphate, 100 mM NaF) and protease inhibitor cocktail (Roche Applied Science). Protein extracts were incubated with each antibody with rotation for 2 hr and then added to protein A/G agarose beads (Santa Cruz, sc-2003) for an additional 1 hr. The immunoprecipitates were washed three times with IP buffer and then resuspended in sample buffer for immunoblotting.

For immunoblotting, protein extracts were prepared in lysis buffer as described before ^54^. Briefly, cells were lysed in lysis buffer (20 mM HEPES, 1% TX-100, 150 mM NaCl, 1 mM EGTA, 1 mM EDTA) containing phosphatase inhibitors (10 mM sodium pyrophosphate, 20 mM β- glycerophosphate, 100 mM NaF, 5 mM IAA, 20 nM okadaic acid (OA)) and protease inhibitor cocktail. Solubilized proteins were collected by centrifugation and quantified using protein assay reagent (Bio-Rad). Samples containing equal amounts of protein were resolved by electrophoresis on a 8-10% gel and transferred to polyvinylidene difluoride membranes (Bio-Rad). Standard enhanced chemiluminescence (ECL) was used for film exposure, and an Azure or LI-COR machine was used to image the blot. ImageJ or LI-COR imaging software was used to quantify bands on the blots. To determine total O-GlcNAc levels, cells were lysed in RIPPA buffer with a Pierce^TM^ protease inhibitor mini tablet and a Pierce^TM^ phosphatase inhibitor mini tablet (1 tablet per 10 mL of lysis buffer). The lysates were incubated on ice for 30 min and centrifuged at 13000 rpm for 10 min before immunoblotting.

### Immunofluorescence

Cells were plated in triplicate in 4 well culture slides. Wild-type or 6SA GFP-tagged SNAIL plasmid (0.75 μg) was transiently transfected by using Lipofectamine 2000 overnight. Cells were fixed in 4% paraformaldehyde (PFA) for 30 minutes followed by four washes in ice cold PBS. After fixation of the cells, the localization of SNAIL proteins (green) and nuclei (blue; Hoechst 33342) were examined under a confocal microscope (Zeiss Lsm 700), in at least 40 randomly selected fields at X400 magnification.

### Hexokinase activity assay

Hexokinase activity in whole-cell lysates and mitochondrial fractions was measured using a standard glucose-6-phosphate (G6P) dehydrogenase (G6PDH)-coupled spectrophotometric assay as described previously ^55^. Whole-cell lysates were prepared by brief sonication in homogenization buffer containing 45 mM Tris-HCl, 50 mM KH2PO4, 10 mM glucose, and 0.5 mM EGTA. Hexokinase activity was measured as the total glucose-phosphorylating capacity of whole-cell lysates in a final assay mixture containing 50 mM triethanolamine chloride, 7.5 mM MgCl2, 0.5 mM EGTA, 11 mM monothioglycerol, 4 mM glucose, 6.6 mM ATP, 0.5 mg/ml NADP, and 0.5 U/ml G6PDH, pH 8.5. Hexokinase activity in each sample was calculated as the coupled rate of NADPH formation by the Lambert-Beer law as follows: [(A340/t)] × dilution factor/[protein], where (6.22 mM-^1^cm^-1^) is the extinction coefficient for NADPH at 340 nm (t=time, [protein]=protein concentration).

### Quantitation of G6P in cells by fluorometric assay

An enzymatic fluorometric assay to quantify G6P was performed following the method described previously ^56^ with minor modifications. Briefly, samples were extracted from 5 × 10^6^ cells with MeOH/CHCl_3_ and stored at -80°C. Just prior to the assay, the extracted samples were dissolved in 50 μl of Millipore water, and then 10 μl of each extraction sample were incubated for 30 min at room temperature in a 96-well plate with 90 μl of an assay cocktail containing 50 mM triethanolamine (TEA, pH 7.6), 1.0 mM MgCl2, 100 μM NADP+, 10 μM resazurin, 1.5 U/ml G6PD, and 0.2 U/ml diaphorase). In this assay, G6P is oxidized by G6P dehydrogenase in the presence of NADP+, and stoichiometrically generated NADPH is then amplified by the diaphorase– resazurin system. Fluorescence at 590 nm was measured using excitation at 530 nm. Background fluorescence was corrected by subtracting the value of the blank for each sample, and G6P concentrations were calculated from a standard curve. Fluorescence was measured using a Tecan Infinite M200 PRO plate reader in 96-well black assay plates.

### Protein Stability Assays

For measuring MCL1 protein stability in M15-4 CHO cells and in M15-4 CHO cells expressing HK2 or for measuring endogenous SNAIL protein stability in 4T1 and 4T1shHK2 cells, cells were plated on 6-well plates at a seeding density of 2 x 10^5^ cells/well. Cells were treated with cycloheximide (100 μM) for the indicated time points. For wild-type GFP-tagged or 6SA GFP-tagged SNAIL, 1.5μg of plasmid was transiently transfected by using Lipofectamine 2000 overnight.

### Enzyme-linked Immunosorbent Assay for Monitoring the HK2/PRKAR1a and HK2/GSK3b Interaction

Enzyme-linked immunosorbent assays were conducted in Pierce Nickel (His) or Glutathione (GST) 96-well coated plates. The plates were respectively coated with His-PRKAR1a (50nM; Sino Biological) or GST-GSK3b (50nM; Abcam), in coating buffer (phosphate-buffered saline containing 0.5 mM phenylmethanesulfonyl fluoride and 1mM dithiothreitol) and incubated overnight at 4°C. Thereafter, unbound protein was removed by washing the wells three times with 100ul of washing buffer (phosphate-buffered saline containing 0.05% Tween 20) and blocking buffer (coating buffer containing 0.6% skimmed milk powder and 0.05% Tween 20) was added (100 ul, 1 h, room temperature). After removal, monitoring of interactions of PRKAR1a or GSK3b with HK2 was carried out in coating buffer and increasing concentration of Myc-HK2 protein (OriGene; 0-64nM, 40ul/well, 2h, room temperature). For G6P-induced binding inhibition, 30minutes before the end of the incubation, G6P (0-100uM) is added directly to the wells. Unbound protein was removed by washing the wells three times with 100ul of washing buffer and bound Myc-HK2 was detected with monoclonal anti-Myc HRP-conjugated antibody (Abcam; 1:5000 in blocking buffer, 1h, room temperature). Bound PRKAR1a or GSK3b were detected by incubation with rabbit anti- PRKAR1a antibody (Cell Signaling; 1:2000 in blocking buffer) or mouse anti-GSK3b antibody (Millipore; 1:2000 in blocking buffer) and HRP-conjugated anti-rabbit or mouse IgG (1:3000, 1 h, room temperature). For each assay, non-specific binding is also conducted as described in figures.

Each antibody incubation step was followed by washing. The HRP reaction was initiated by addition of 3,3’,5,5’-tetramethylbenzidine enzyme-linked immunosorbent assay substrate solution (Sigma) and terminated after 30 min by adding H2SO4 (2M; 50ul/well). The colored reaction product was quantified by measuring A450 nm in a Cytation 1 plate reader (Biotek).

### Enzyme-linked Immunosorbent Assay for Monitoring the PRKAR1a/HK2/GSK3b Interaction

Binding assay is conducted on Pierce Nickel-coated 96 well plates and after successive incubation with His-PRKAR1a and Myc-HK2 as described above, unbound proteins will be washed and blocking buffer will be added for 1h at room temperature. Thereafter, GST-GSK3b is added at increasing concentration (0-64nM; 40ul/well, 2h, room temperature). For G6P-induced binding inhibition, 30minutes before the end of the incubation, G6P (0-1000uM) is added directly to the wells. Unbound protein was removed by washing the wells as described above and bound GST- GSK3b was detected with monoclonal anti-GST HRP-conjugated antibody (Abcam; 1:5000 in blocking buffer, 1h, room temperature). Bound PRKAR1a or HK2 were detected by incubation with rabbit anti- PRKAR1a antibody (Cell Signaling; 1:2000 in blocking buffer) or rabbit anti-HK2 antibody (Cell Signaling; 1:2000 in blocking buffer) and HRP-conjugated anti-rabbit IgG (1:3000, 1 h, room temperature). For each assay, non-specific binding is also conducted as described in figures and HRP reaction is conducted as described above.

### Monitoring of GSK3 phosphorylation in the HK2/PRKAR1α/GSK3β complex

Enzyme-linked immunosorbent assays were conducted in Pierce Nickel- 8-well-strip coated plates. The plates were coated with His-HK2 (50nM; Creative Biomart) in coating buffer and incubated overnight at 4°C. Unbound proteins is then washed and blocking buffer is added for 1h at room temperature. Thereafter, Myc-PRKAR1α (32nM; Creative Biomart), PKAc (20nM; Promega) and GST-GSK3β (32nM; Abcam) successively, each for 1h incubation at room temperature, and unbound protein will be carefully washed after each incubation. After all proteins are bound, kinase buffer (1mM ATP and 10mM MgCl_2_ in Tris 20mM, pH7.4) with or without cyclic AMP (10µM) is added to all appropriate wells (1h; room temperature). After incubation, kinase buffer is washed and appropriate antibody will be added to the wells de detect bound proteins (P-GSK3β, total GSK3, PRKAR1α, HK2 and PKAc) and incubated overnight at 4°C. After washing, total bound GST-GSK3β was detected with monoclonal anti-GST HRP-conjugated antibody, and bound P-GSK3β, PRKAR1α, HK2 and PKAc antibodies are detected with HRP-conjugated anti-rabbit IgG (1:3000, 2h, room temperature). The HRP reaction is conducted as described above.

For G6P-induced binding inhibition, G6P (1mM) is added 30minutes before the end of the incubation with GST-GSK3β, before the addition of kinase buffer.

Negative control reactions are conducted as described above, in absence of His-HK2 or Myc- PKAR1α.

### Enzyme-linked Immunosorbent Assay for Monitoring the PRKAR1a/HK2 Interaction in presence of FMP-API-1

Binding assay is conducted on Pierce Nickel-coated 8-well strip plates and after successive incubation with His-PRKAR1α and Myc-HK2 as described above, unbound proteins will be washed and blocking buffer will be added for 1h at room temperature. Thereafter, FMP-API-1 is added directly to the wells 30minutes before the end of the incubation with Myc-HK2 at increasing concentration (0-1000uM). Unbound protein was removed by washing the wells as described above and bound Myc-HK2 was detected with monoclonal anti-Myc HRP-conjugated antibody (Abcam; 1:5000 in blocking buffer, 1h, room temperature). HRP reaction is then conducted as described above.

### Metabolic labelling

For metabolic tracing in A549 cells, 7.0 × 10^5^ cells were plated onto 6 well plates for next day pulse-labeling with either 25 mM [1,2-^13^C] glucose (Cambridge Isotope Laboratories). Isotope labeling experiments were performed for 4 hours and extra plates were included for cell counts. At the time of collection, plates were washed twice with 1 mL of (9 g/L) NaCl. Then, cells were incubated with 600 μL mixed solvent (water:methanol:acetonitrile = 1:1:1) containing 2 μL of 2 mg/mL Norvaline (SigmaAldrich) dissolved in distilled water as an internal standard, and then scraped down with cell scrapers. The solution was shaken at 1,200 rpm for 30 min and centrifuged at 16,000 × g for 15 min at 4 °C. The supernatant (280 μL) was transferred to a clean tube and lyophilized under nitrogen gas. The lyophilized samples were derivatized with 15 µL of 2 wt% methoxylamine hydrochloride (Thermo Fisher) for 60 min at 42 °C. Next, 35 µL of N-methyl-N-(tert-butyldimethylsilyl)-trifluoroacetamide (MTBSTFA) + 1% tert- butyldimetheylchlorosilane (t-BDMCS) (SigmaAldrich) was added and the samples were incubated for 60 min at 75 °C. The derivatized samples were then centrifuged at 16,000 × g for 5 min at 4 °C, and the supernatant (1 μL) was subjected to GC-MS measurement.

GC-MS analysis was performed using an Agilent 7890B GC equipped with a DB-5ms column (30 m × 0.25-mm inner diameter; film thickness, 0.25 μm; Agilent J&W Scientific). The front inlet temperature was 280 °C and helium flow was maintained at 1.428 mL/min. For intracellular lactate analysis, the column temperature was held at 60 °C for 1 minute followed by 1-minute of run time, rising at 10 °C/minute to 320 °C and holding for 1 minutes followed by a post run of 320 °C for 9 minutes. For the analysis of G6P and 6PG, the initial rate was held at 60 °C for 1 minute followed by 1-minute of run time, rising at 10 °C/minute to 325 °C and holding for 10 minutes followed by a post run of 60 °C for 1 minutes ^57, 58^. The peak area of each quantified ion was calculated and corrected for natural isotope abundances by following the procedure by Fernandez et al ^59^, and then normalized by the peak area of norvaline as an internal standard and cell number.

For metabolic tracing, in 4T1 and 4T1shHK2 cells, the cells were plated on 6cm plate in triplicate (3 x 10^5^) in 5mM glucose DMEM media for 48 hours. Cells were washed in PBS and 5mM of uniformly labeled C-13 glucose in DMEM with 10% dialyzed FBS was added for 5 minutes following extraction in 1mL of cold 90/10 acetonitrile/water. Cells were collected using a cell scrapper, vortexed for 2 minutes, and centrifuged at highest speed for 5 minutes. The supernatant was transferred to clean eppendorf tube and sent to MD Anderson Cancer Center’s Proteomics and Metabolomics Core Facility for analysis by Ion chromatography mass spectrometry (IC-MS). One 6cm plate was collected for western blot. Cell number was counted on extra 6 well plate and protein concentration was quantified with remaining cell pellet.

### Extracellular lactate measurements

Extracellular lactate concentrations were measured using a YSI 2700 Bioanalyzer (YSI).

### Mouse strains and treatment protocol

C57BL/6 *MMTV-PyMT* and *HK2^f/f^;UBC-Cre^ERT2^* mice were previously described ^35^. *MMTV-PyMT;HK2^f/f^* mice were crossed with *HK2^f/f^;UBC-Cre^ERT2^* or *HK2^f/^*^f^ mice to generate experimental and control mice on a C57BL/6 background. Once a primary tumor was palpable (∼12 weeks old), 0.1 ml of 30 mg/ml tamoxifen was injected IP for 7 consecutive days to systemically delete HK2 as previously described ^35^. At the tumor end point (20% weight loss from baseline, 15% weight gain compared to aged-matched controls, tumor size greater than 15% of the body weight, tumor mass > 2 cm, tumor ulceration, pallor, respiratory distress, or inability to ambulate), the mice were euthanized, and their lungs were analyzed for metastases.

MMTV-rtTA and LSL-luc mice were purchased from Jackson Laboratory.

*MMTV-PyMT;Hk2^f/f^;LSL.Luc;MMTV.rtTATet(O)Cre* mice were generated by first crossing *MMTV-rtTA mice with LSL.Luc* mice to generat*e MMTV-rtTA:LSL.Luc mice.* These mice were crossed *with MMTV-PyMT;Hk2^f/f^ mice*.

Balb/cJ mice were purchased from Jackson Laboratory and NOD.Cg-Prkdcscid (NOD-F) mice from TACONIC.

### Histochemistry

Mammary tumor tissues and PBS-inflated lung lobes were collected, and macroscopic lung lesions were counted visually. The tissues were fixed in 10% formalin for 48 hr. Tissues were paraffin embedded and sectioned for hematoxylin and eosin (H&E) staining (5-µm sections). Microscopic lung lesions were counted using a microscope.

### Primary tumor isolation and processing

Tumors isolated from MMTV-PyMT mice were isolated for scRNA-seq, western blotting, and orthotopic transplantation. The mice were sacrificed close to the tumor end point (primary tumor >2 cm). Primary tumors were dissected and washed several times in PBS with 1× pen/strep. The tumors were chopped into 1-mm pieces with a sterile scalpel and placed in collagenase for 1 hr at 37°C on a shaker. The collagenase mixture consisted of 10% type IV collagenase and 1% DNase dissolved and sterile filtered in DMEM. After digestion, the cells were spun for 5 min at 1000 rpm. The supernatant was aspirated, and 2 ml of ACK lysis buffer was used to resuspend the pellet for 2 min. The mixture was centrifuged again, and the pellet was washed in DMEM or PBS at least twice before proceeding with the experiment. For scRNA-seq, the pellet was resuspended in PBS and filtered through a 40-µm filter to isolate single cells for sequencing. The cells were counted with trypan blue staining to determine cell viability before proceeding with the Drop-seq protocol as previously described ^39–41, 60^.

### Single Cell RNA-seq and bioinformatics

For scRNA-seq, the PyMT tumors were isolated as described in the section regarding primary tumor isolation and processing. The pellet was resuspended in PBS and filtered through a 40-µm filter to isolate single cells for sequencing. The cells were counted with trypan blue staining to determine cell viability before proceeding with the Drop-seq protocol ^38^. To isolate single-cell droplets, 1.65 × 10^5^ cells were resuspended in 1.5 ml of 0.1% BSA PBA and loaded on a microfluidic chip. Cells were lysed in the droplet, and mRNA was bound to a unique molecular identifier (UMI) on barcoded beads. The individual droplets were broken down and pooled in a reverse transcriptase mixture. Single-cell transcriptomes attached to microparticles (STAMPs) were created by reverse transcription. The STAMPS were PCR amplified, and unhybridized DNA was removed with exonuclease I treatment. The UIC Research Resources Center (RRC) used TapeStation and Qubit to check the cDNA quality and quantity of the amplified products. The Nextera XT kit was used to make the libraries, which were then sequenced by an Illumina NextSeq 500. The raw sequence data were filtered and aligned to the mouse genome (mm10) with the addition of the PyMT gene sequence to help identify positive tumor cells. The digital expression matrix file containing UMIs were analyzed with the Seurat package version 2.3.4 ^61^ R version 3.5.3, and clusters were grouped based on similar gene expression.

### Quantitative RT-PCR

Total RNA was extracted with TRIzol reagent (Invitrogen) or a Qiagen RNeasy mini kit. Quantitative PCR was performed with a BIO-RAD iTaq Universal SYBR Green One-Step Kit and system. Each sample was prepared in triplicate and normalized to B-actin mRNA levels.

### Orthotopic transplantation

Cells from *MMTV-PyMT;HK2^f/f^* mice were isolated as described above. The pellet was resuspended in PBS and filtered through a 75-µm filter before the cells were plated in 10% FBS, 1% pen/strep, and high-glucose DMEM. The cells were then infected with either adenovirus expressing GFP or adenovirus expressing GFP-Cre at a MOI of 1000 to delete HK2. Deletion of HK2 was verified via western blotting. The cells were orthotopically transplanted into the mammary fat pads of NOG-F mice. Briefly, the cells were trypsinized, resuspended, and counted with trypan blue staining to determine cell viability and number. Cells (1 x 10^5^) were resuspended in 100 µl of a 1:1 mixture of PBS and Matrigel (Corning). They were then transplanted into the fourth mammary fat pads of NOG-F mice. Similarly, approximately 5 × 10^5^ 4T1 and 4T1shHK2 cells resuspended in 100 µl of a 1:1 mixture of PBS and Matrigel (Corning) were transplanted into the fourth mammary fat pads of syngeneic Balb/cJ mice. The recipient mice were monitored until they reached the tumor end point. The lungs were then isolated and processed for H&E staining to quantify lung metastasis. Finally, 5 x 10^4^ cells of 4T1shHK2 with dox inducible Empty Vector or 6SA-SNAIL cells were resuspended in 100ul of 1:1 in PBS and Matrigel (Corning) mixture and transplanted into the fourth mammary fat pad of syngeneic Balb/cJ mice. The following day DOX (1ml/ml in water with 5% sucrose) was added to both groups of mice for 1 week. The recipient mice were monitored until they reached tumor end-point and lungs were isolated and processed for H&E stain to quantify lung metastasis.

### Transwell migration and invasion assays

For the transwell migration assay, 1 × 10^5^ cells in 200 µl were incubated in 24-plate chamber wells containing a 8.0-μm filter in high-glucose DMEM with 0% serum. The lower chamber contained 750 µl of high-glucose DMEM with 20% serum as a stimulant. After 12 hr, the cells were washed twice in PBS, and the nonmigrated cells were scraped away with a cotton swab. The cells were fixed in methanol and stained with crystal violet. Migrated cells in five random fields were counted with ImageJ. For the transwell invasion assay, 1 × 10^5^ cells in 500 µl were incubated in gel-coated 24-plate chamber wells containing a 8.0-μm filter in DMEM containing 5 mM glucose and 0% serum. The lower chamber contained 750 µl of DMEM containing 5 mM glucose and 20% serum as a stimulant. After 24 hr, the cells were washed twice in PBS, and the nonmigrated cells were scraped away with a cotton swab. The cells were fixed with methanol and stained with crystal violet. Migrated and invaded cells in ten random fields were counted with ImageJ.

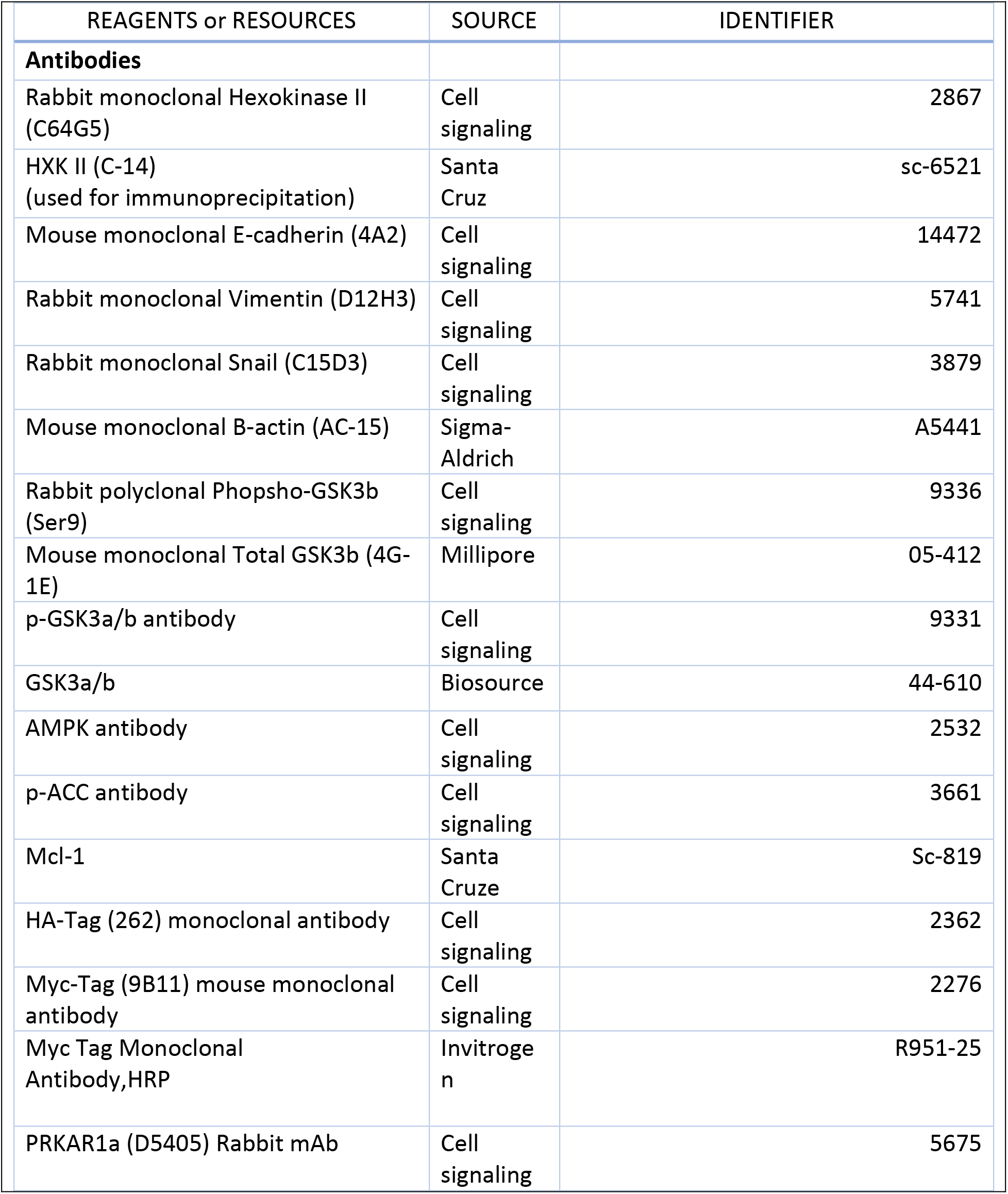
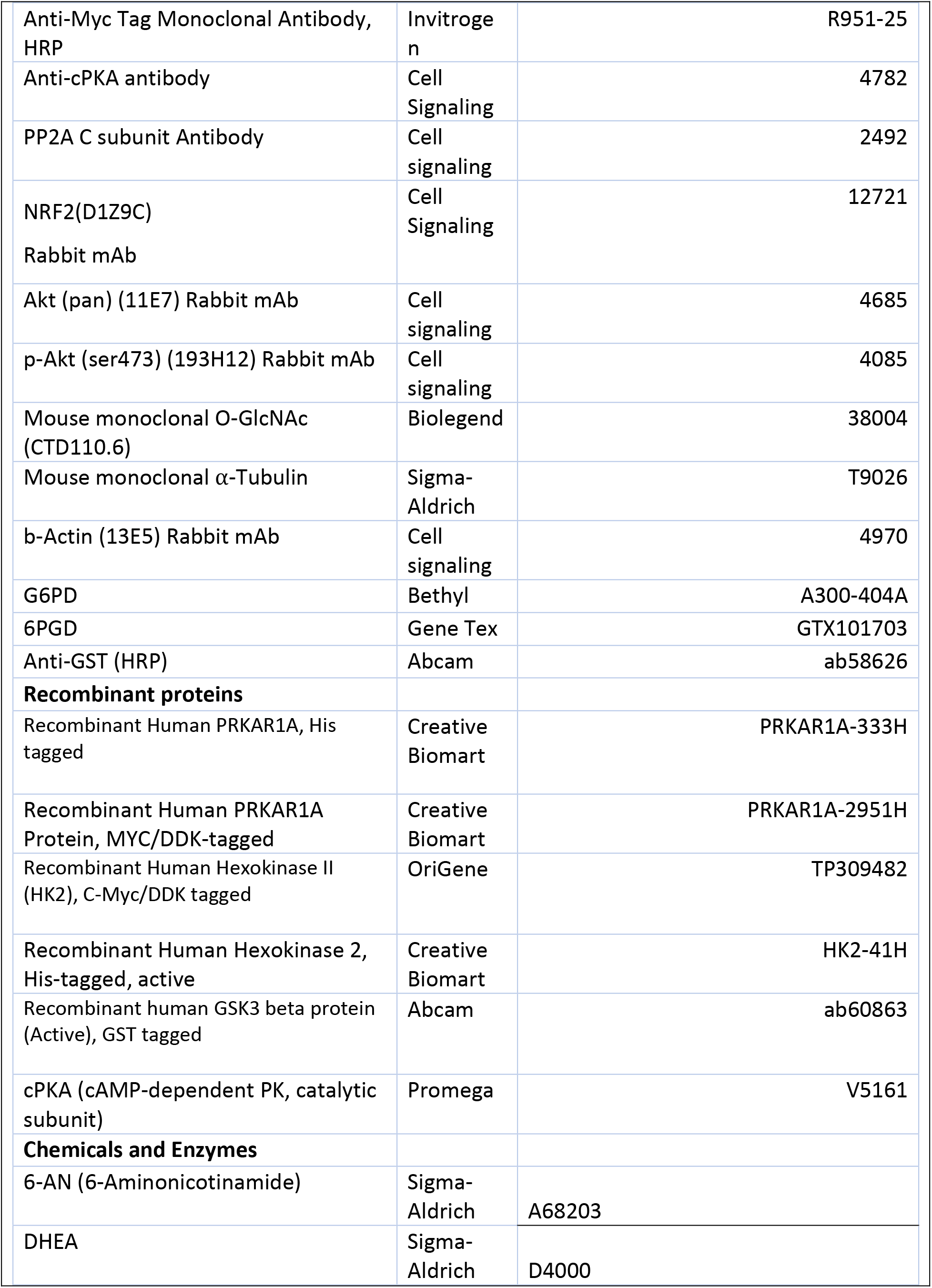
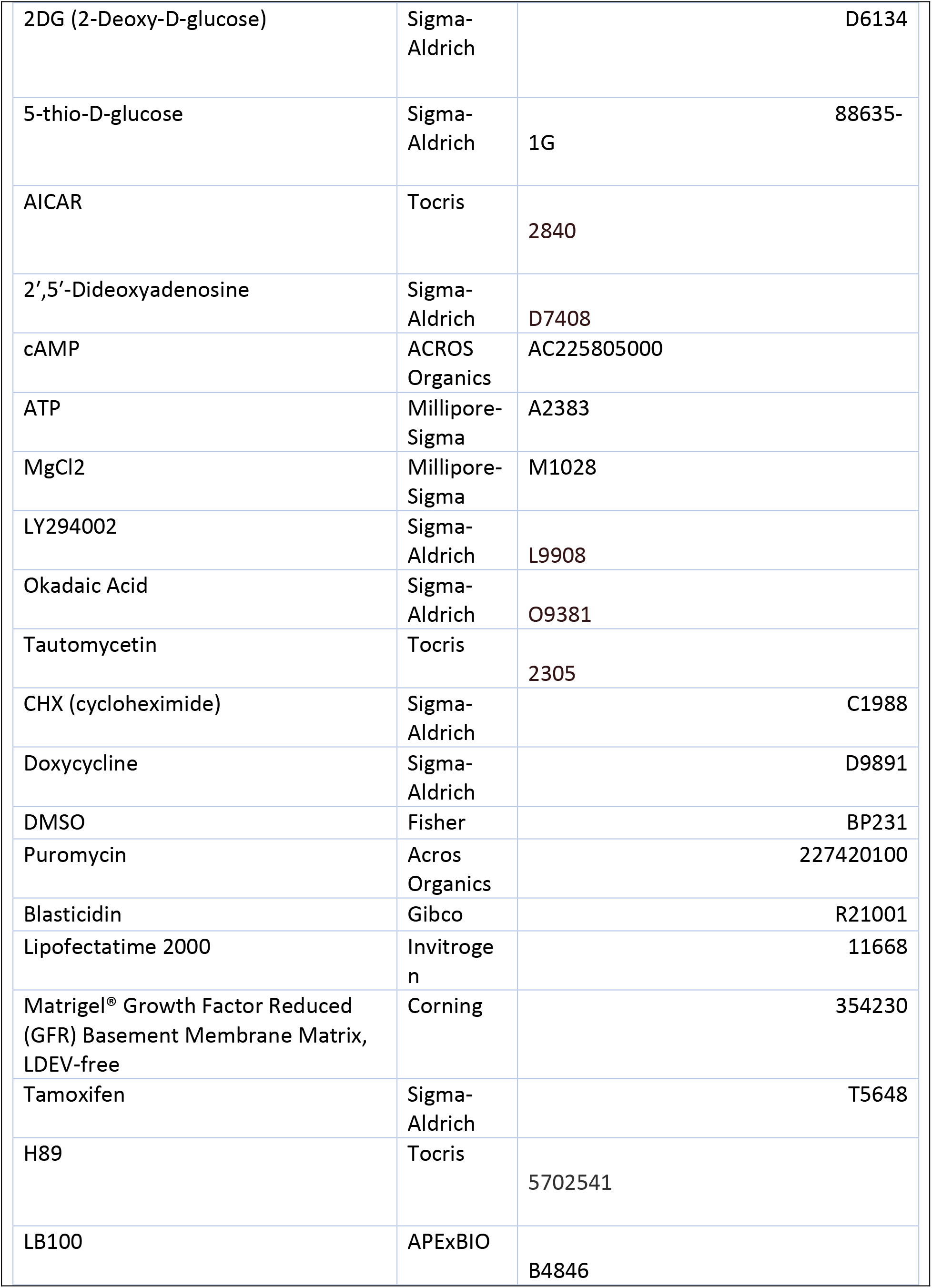
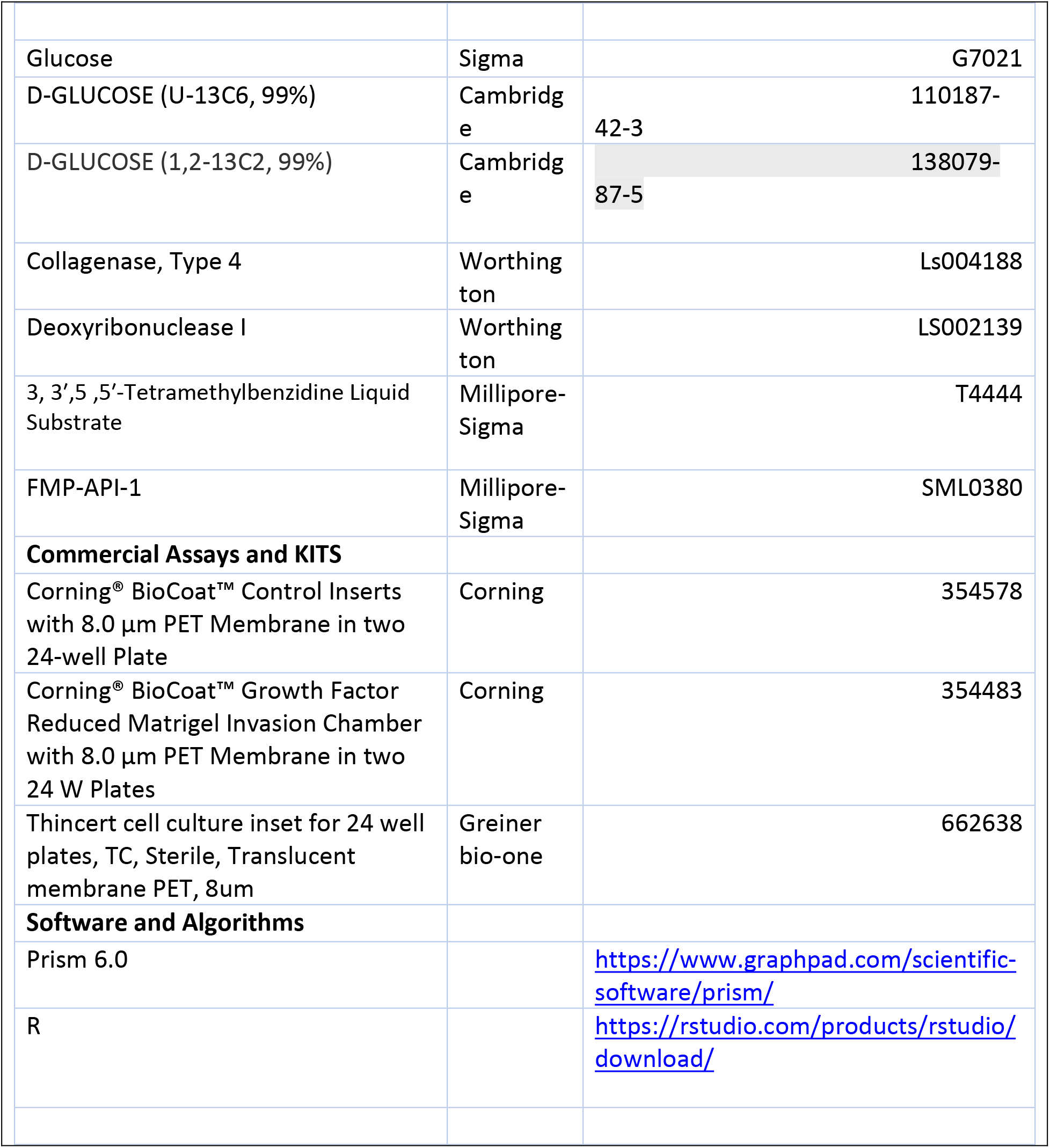
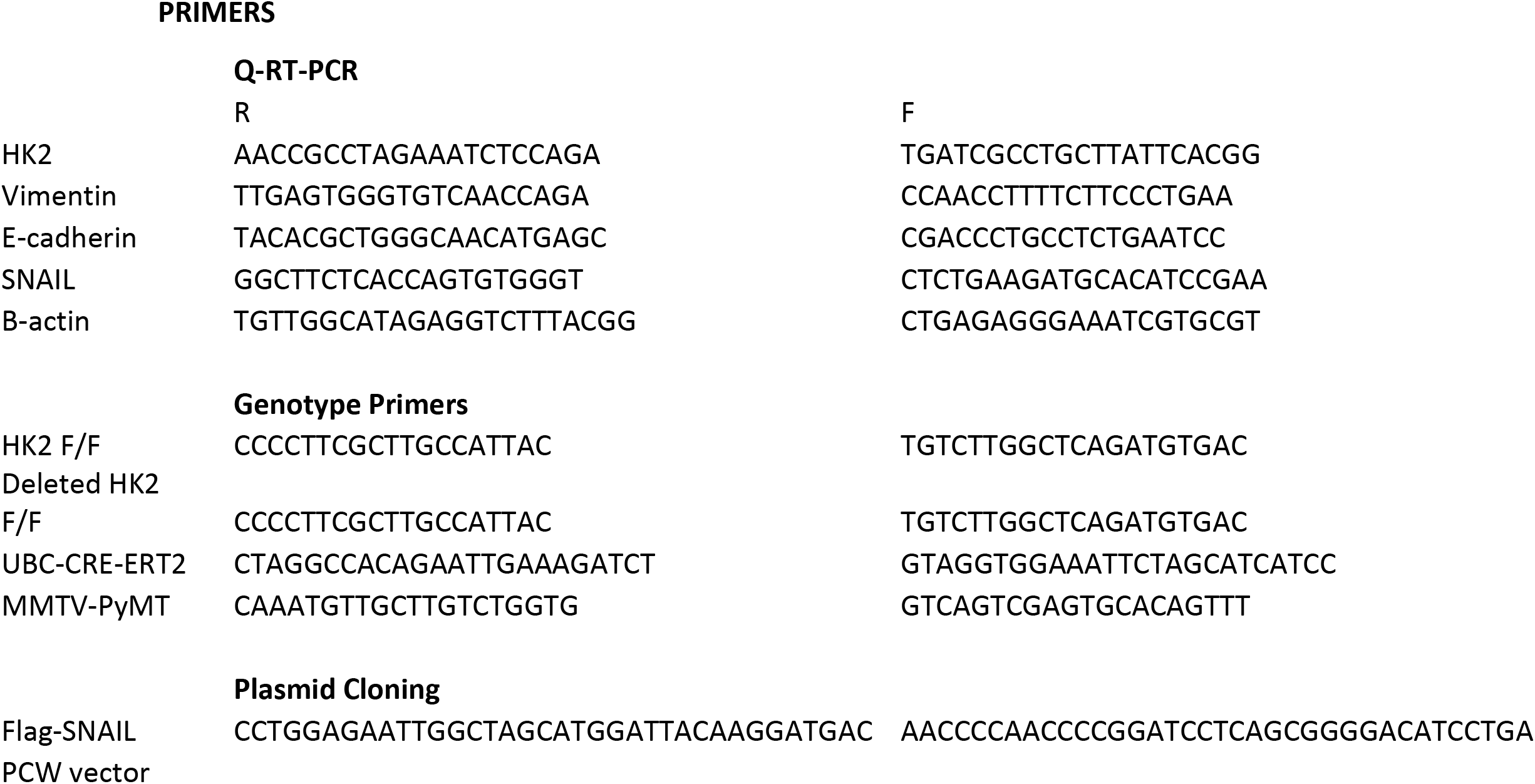

## Acknowledgments

N.H. acknowledges the support from NIH grants R01AG016927, R01CA090764, and R01 CA206167, the VA merit award BX000733, and the VA research career scientist award IK6BX004602. C.B acknowledges the support from F30CA228191. H.R. acknowledges the support from the center for clinical and translational science. A.R.T. acknowledges the support from F30CA225058. M.V.F. acknowledges the support from NIH grant R35GM131707. We would like to thank Ling Jin for maintaining and genotyping the mice.

## Extended Data Figure Legends

**Extended Data Figure 1:**
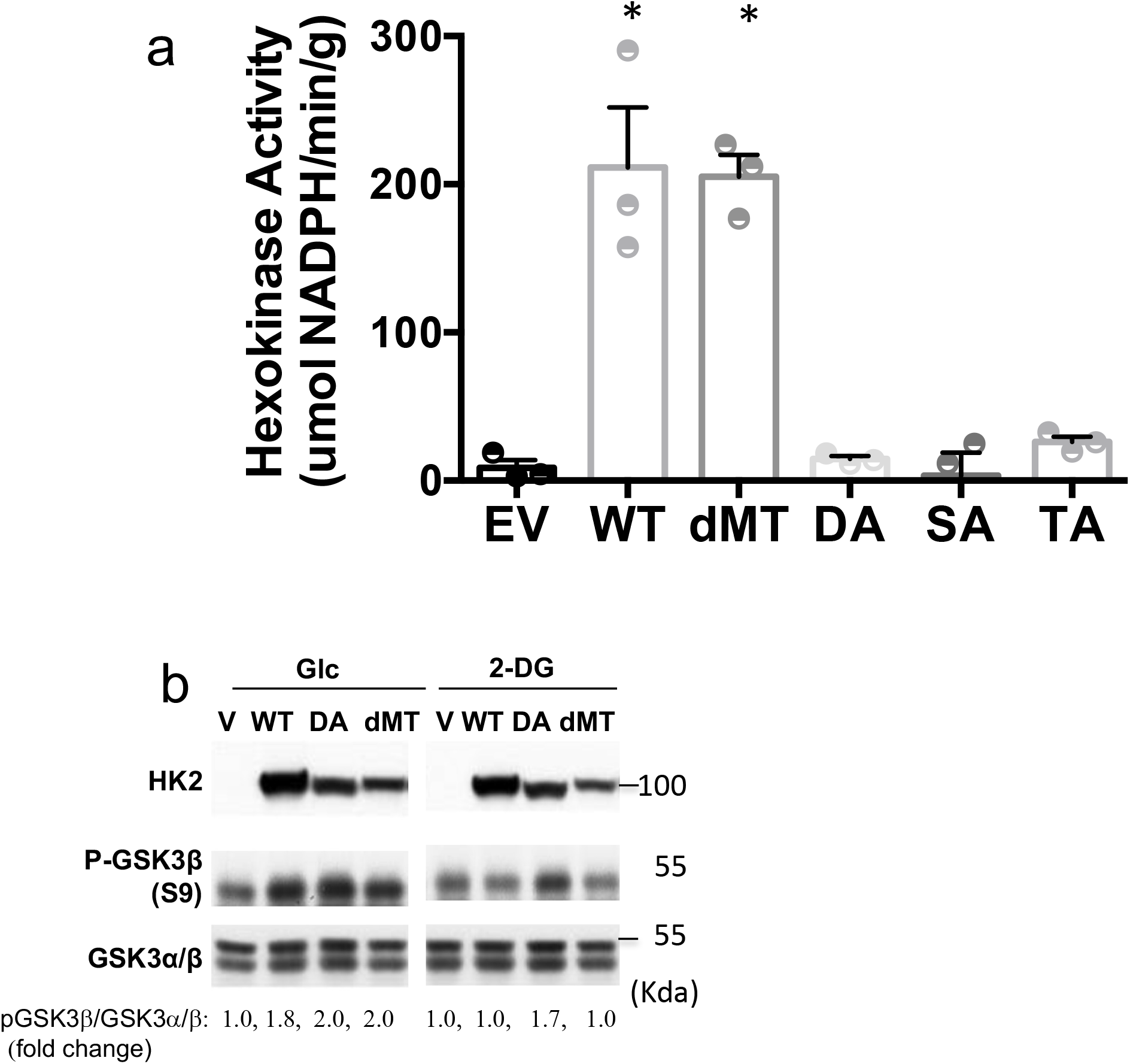
**a.** Hexokinase activity in M15-4 CHO cells expressing empty vector (EV), WT HK2 or HK2 mutants. Results are the mean ± SEM of 3 independent experiments in triplicate. *p<0.05, all 2-sided t-test vs. EV. **b.** MI5-4 CHO cells expressing either wild type (WT), kinase-dead HK2 mutant (DA), mitochondrial binding deficient mutant (dMT) or empty vector (V) were incubated in glucose free medium in the presence of 10mM glucose (G) or 2-DG (D). After 2hr, cells were harvested and analyzed by immunoblotting.

**Extended Data Figure 2:**
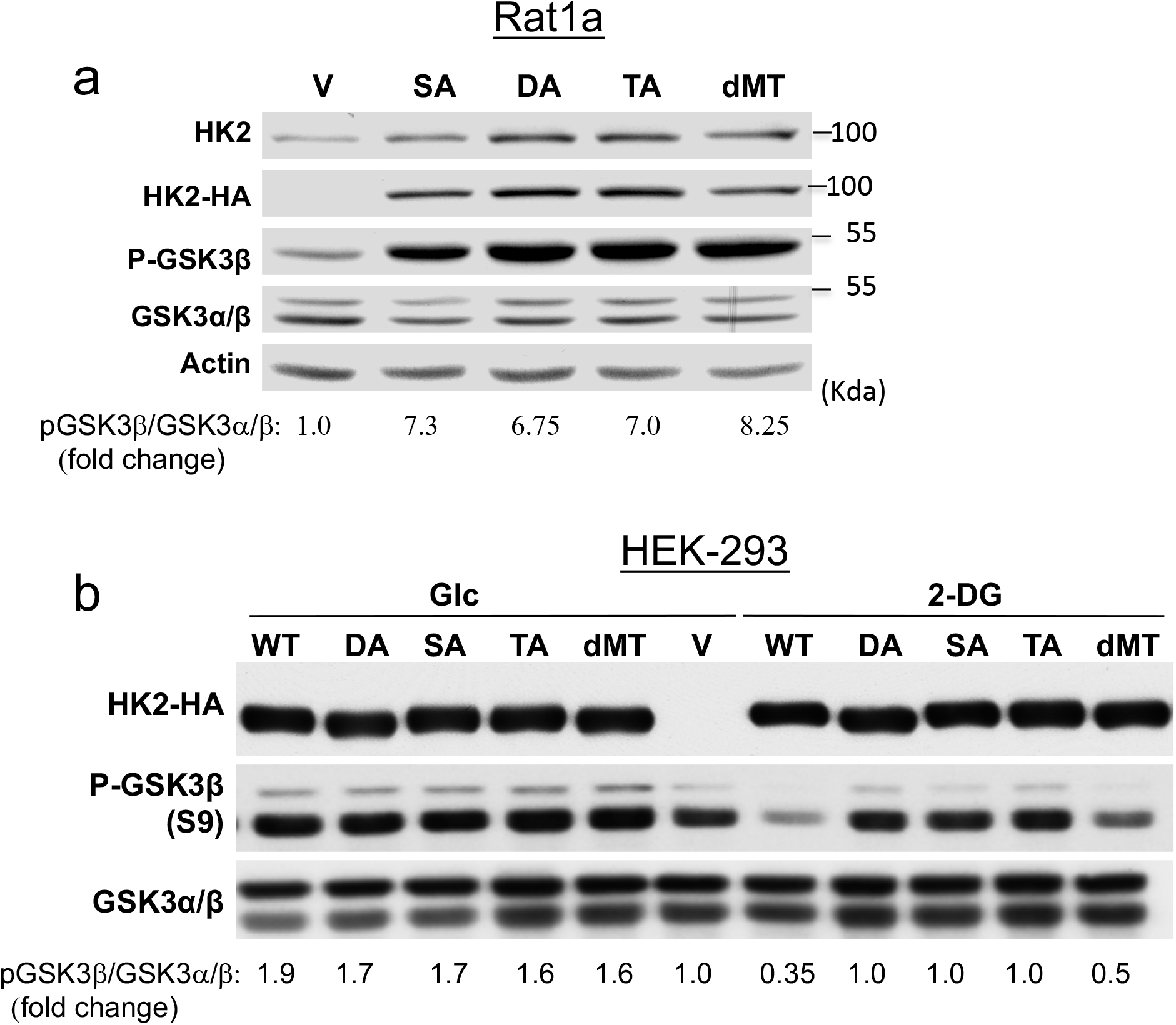
Hexokinase maintains GSK3β phosphorylation independent of its activity, but its activity is required to suppress GSK3β phosphorylation by 2-DG. **a.** The effect of WT HK2 and HK2 mutants overexpression on GSK3β phosphorylation in Rat1a cells. **b.** The effect of 2-DG on GSK3β phosphorylation mediated by either WT HK2 or HK2 mutants in HEK293 cells.

**Extended Data Figure 3:**
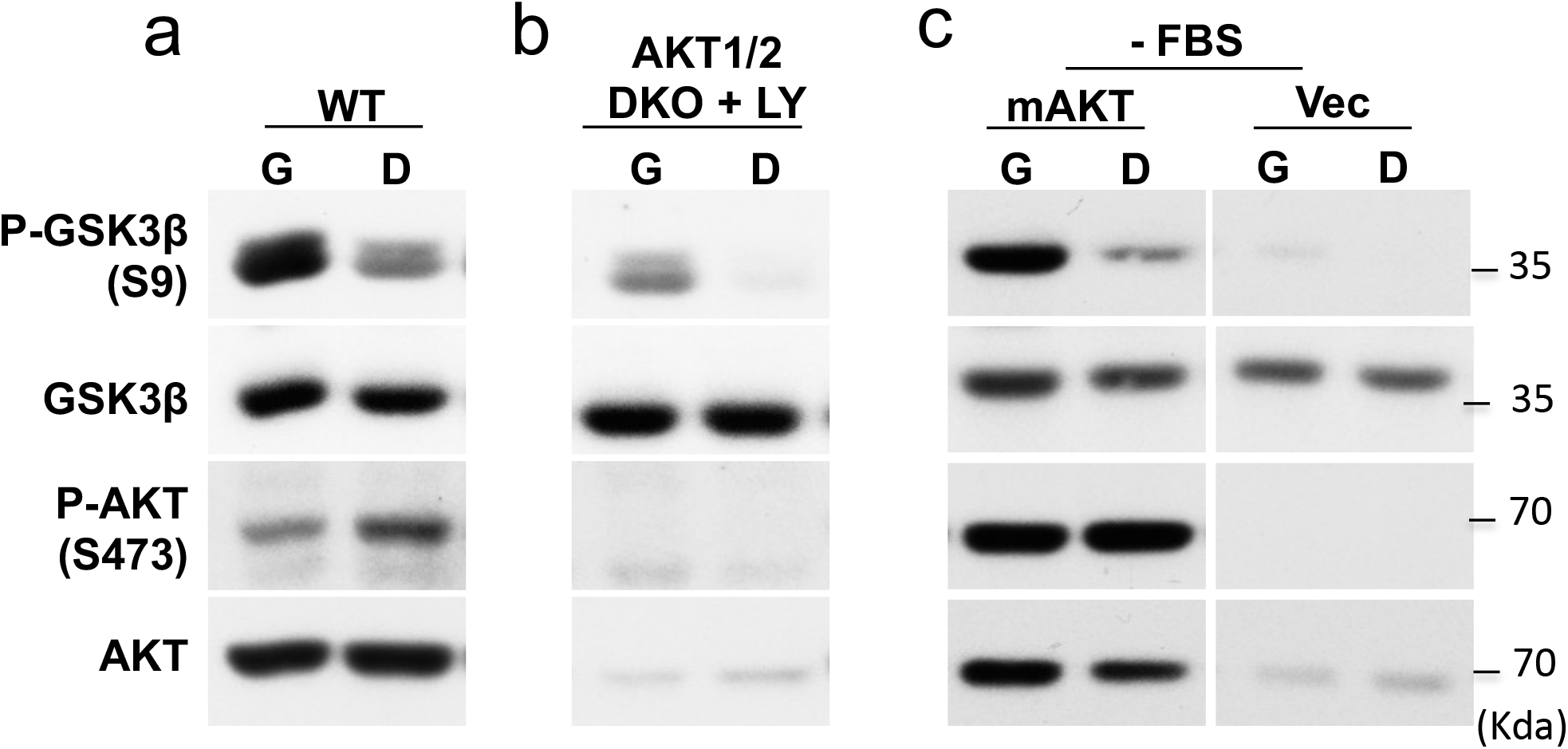
The effect of HK2 on GSK3β phosphorylation is independent of Akt activity. **a.** WT MEFs were incubated in glucose free medium in the presence of 10mM glucose (G) or 2- DG (D) followed by immunoblotting to determine GSK3β and Akt phosphorylation. **b.** Akt1/2 DKO MEFs treated with LY294002 (LY) were incubated in glucose free medium in the presence of 10mM glucose (G) or 2-DG (D) followed by immunoblotting to determine GSK3β and Akt phosphorylation. **c.** MEFs expressing mAkt or vector control (Vec) were deprived of FBS and incubated in glucose free medium in the presence of 10mM glucose (G) or 2-DG (D) followed by immunoblotting to determine GSK3β and Akt phosphorylation.

**Extended Data Figure 4:**
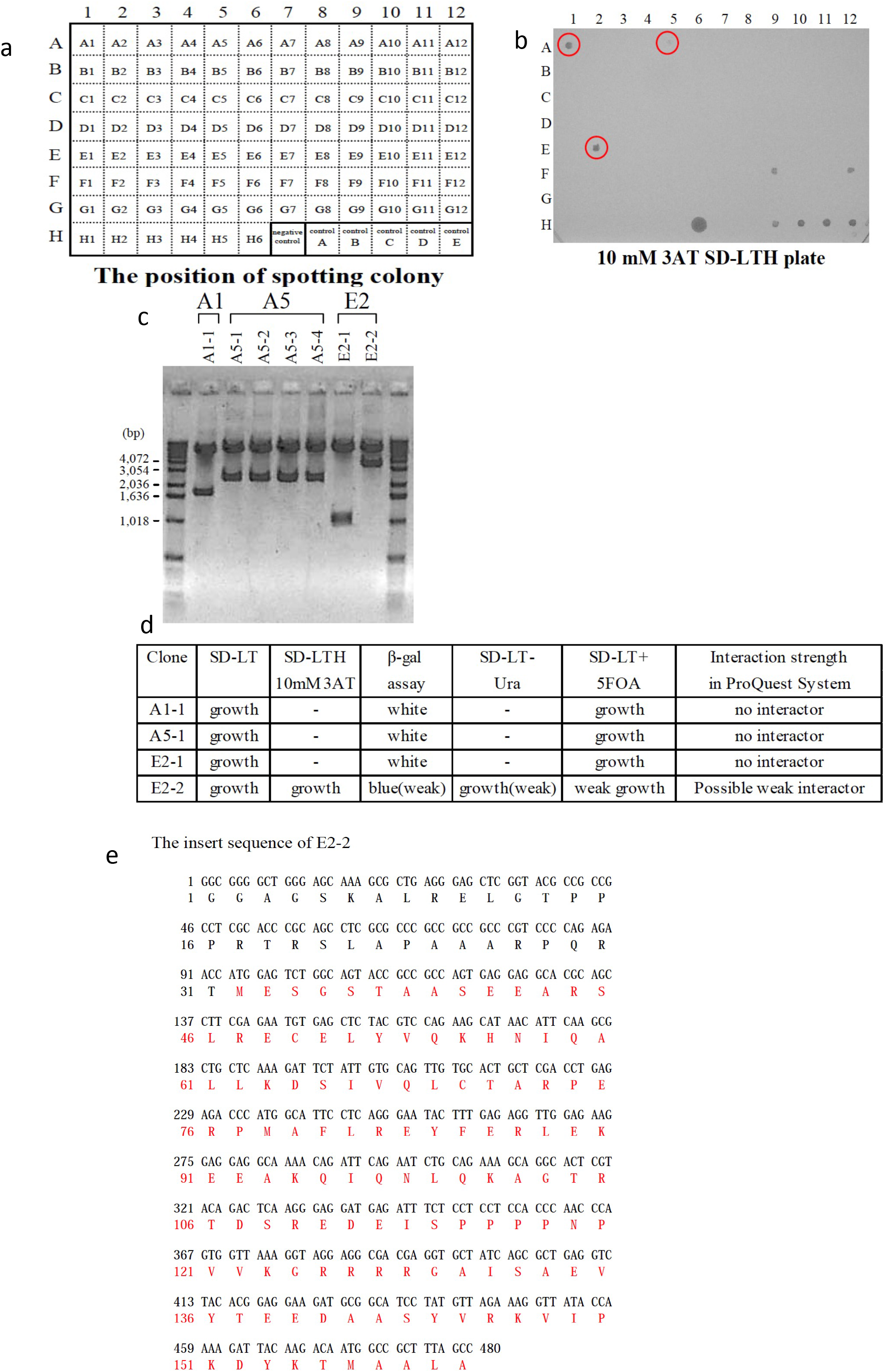
Yeast two-hybrid screen for HK2 interacting proteins. (Performed by Invitrogen life technologies, Japan). Full length HK2 was cloned into the pDEST32 vector as a plasmid bait. Empty pDEST32 or pDEST32 expressing HK2 were transformed into MaV203 yeast competent cells. Large scale screening was performed under 10 mM 3AT concentration. After 4 days incubation, comparatively large 90 colonies were selected and cultured in 100 ul of SD-LTH medium with 10mM 3AT for one day using a 96-well plate, and then spotted on SD-LT plate, SD-LTH 10 mM 3AT plate, and nylon membrane on YPD plate for beta-gal assay. Prey plasmids, purified from possible positive clone, were introduced into E.coli and were estimated fragment size by colony PCR. Plasmids then were purified from the E. coli and transformed along with the bait or empty plasmid back into yeast and tested for all four phenotypes (sensitivity to 3AT, growth on medium without uracil, sensitivity to 5-FOA and detection of β-galactosidase activity). The insert of potential interactor was sequenced and then BLAST search was performed. **a.** Position of spotting clones. **b.** Possible positive clones (grown on 10mM 3AT plates); A1, A5, and E2 (F9,F12, and H6 were false positive. **c.** Inserts in A1, A5 and E2 were amplified by PCR and cloned into plasmids. Insert size after cloning is shown. **d.** Plasmids were re-validated for interaction with the HK2 bait and one plasmid E2-2 was found as a real interactor. **e.** The insert sequence of E2-2. Red labeled amino acids.

**Extended Data Figure 5:**
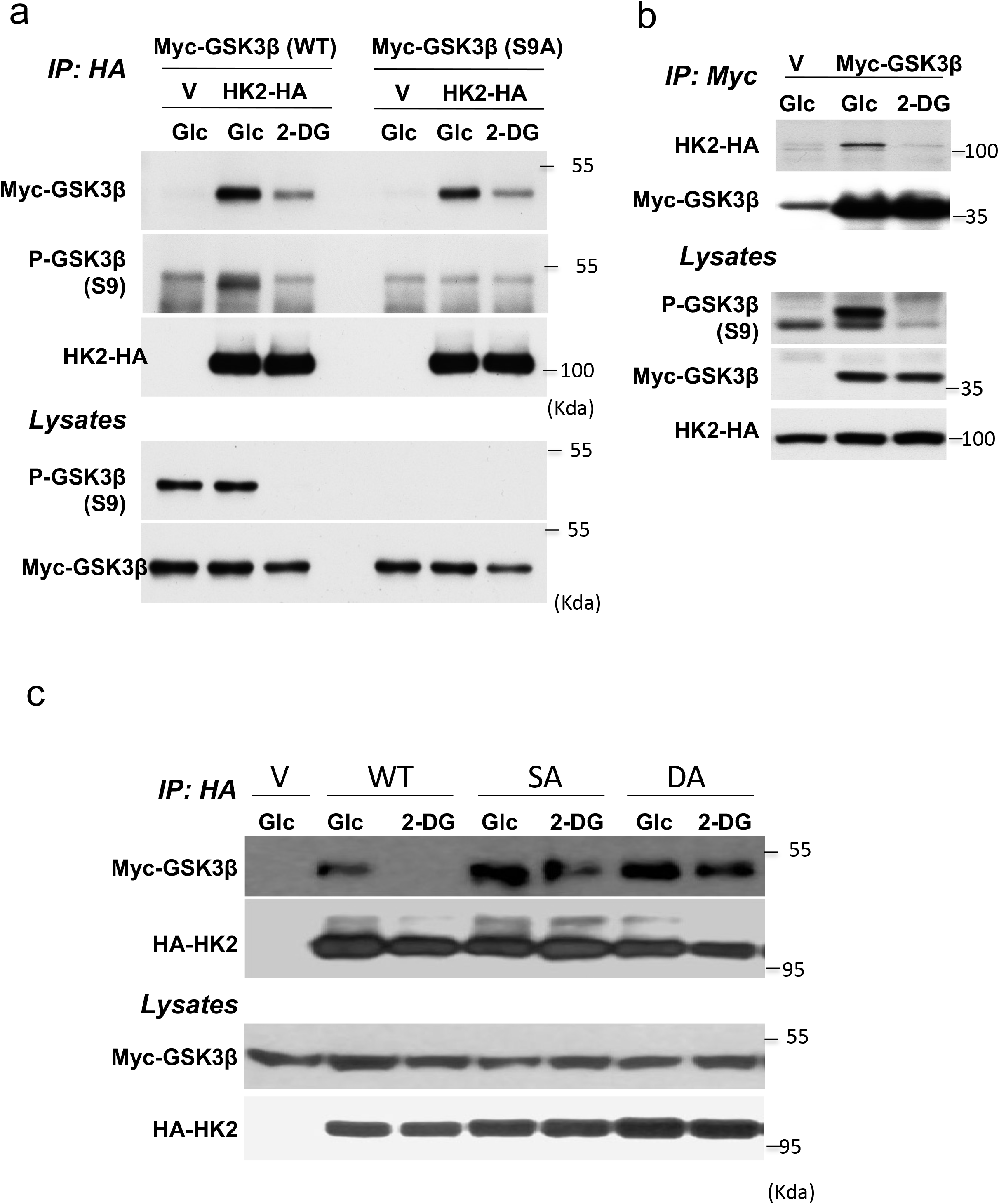
HK2 interacts with GSK3β in a 2DG-dependent manner. **a.** After co-transfection of HK2-HA or HA-vector with Myc-GSK3β wild type (WT) or Myc-GSK3β nonphosphorylatable mutant (S9A) into HEK293 cells, the cells were incubated in glucose free medium in the presence of 10mM glucose (Glc) or 2-DG. After 2hr, cells were lysed for immunoprecipitation with anti-HA antibody followed by immunoblotting using anti-Myc-HRP, anti-HA and anti-P-GSK3β antibodies. Total lysates were subjected to immunoblotting using anti-P-GSK3β, and anti-Myc-HRP antibodies. **b.** After transfection of control Myc-vector or Myc-GSK3β plasmid into HEK293-HK2-HA expressing cells, the cells were incubated in glucose free medium in the presence of 10mM glucose (Glc) or 2-DG. After 2hr, cells were lysed for immunoprecipitation with anti-Myc antibody followed by immunoblotting using anti-HA and anti-Myc-HRP, antibodies. Total lysates were subjected to immunoblotting using anti-P-GSK3β, anti-Myc-HRP, and anti-HA antibodies. **c.** After transfection of Myc-GSK3β into M15-4 CHO cells expressing either WT or mutants HK2, cells were incubated in glucose free medium in the presence of 10mM glucose (Glc) or 2-DG. After 2hr, cells were lysed for immunoprecipitation with anti-HA antibody followed by immunoblotting using anti-Myc-HRP and anti-HA and antibodies. Total lysates were subjected to immunoblotting using anti-Myc-HRP and anti-HA antibodies.

**Extended Data Figure 6:**
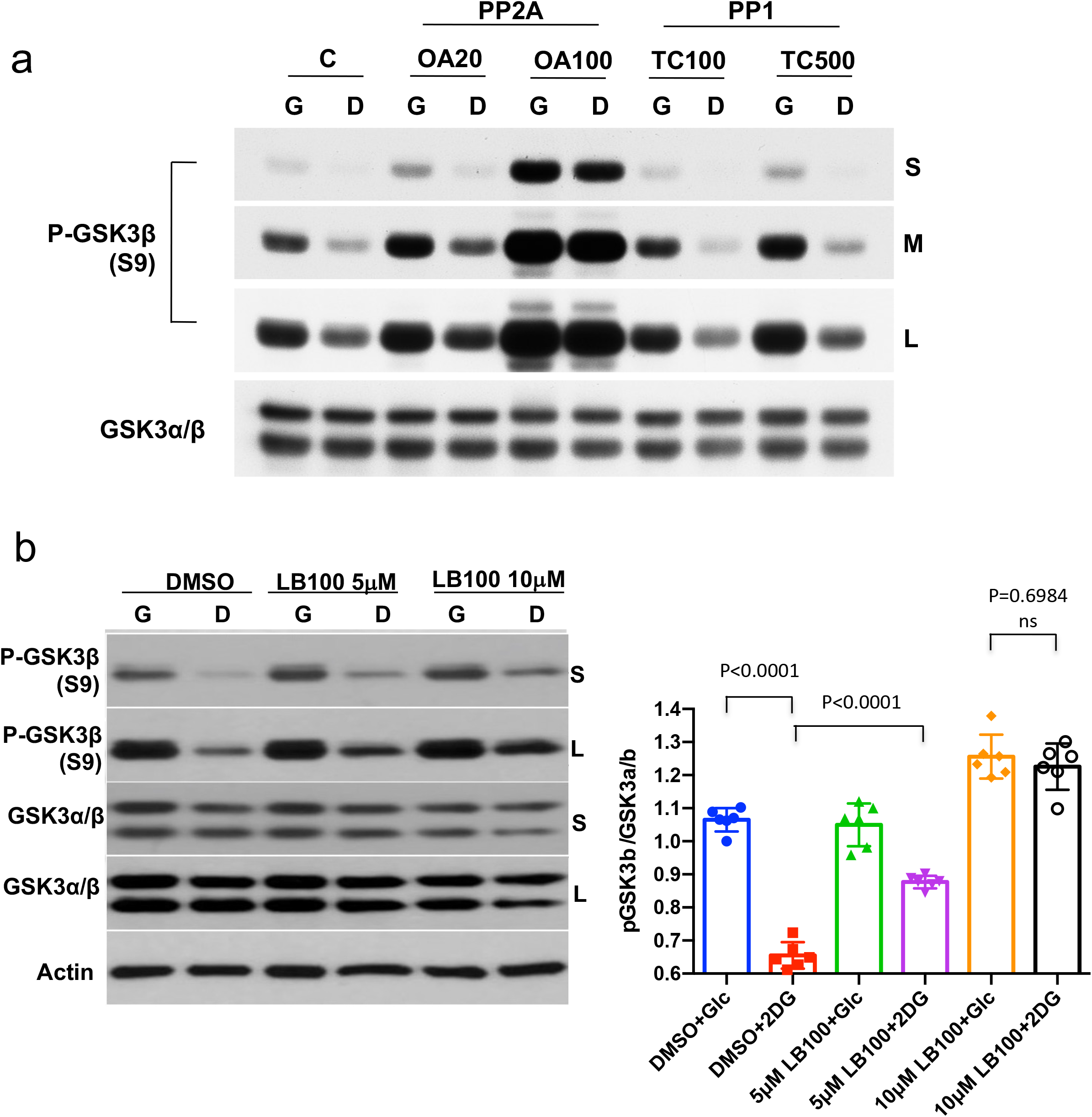
The effect of protein phosphatases on GSK3β phosphorylation. **a.** HeLa cells were incubated in glucose free medium in the presence of 10mM glucose (G) or 2- DG (D). DMSO (C), OA (20nM, 100nM), or TC (100nM, 500nM) were also treated with glucose or 2DG. After 2hr, cells were harvested and analyzed for immunoblotting using anti-P-GSK3β and anti-GSK3α/β (S-short exposure, M-medium exposure, L-long exposure). **b.** Experiment was done as in a except that cells were treated with LB100 quantification of pGSK3b/GSK3a/b ratio is shown. Results are the mean ± SEM of 3 independent experiments in duplicates.

**Extended Data Figure 7:**
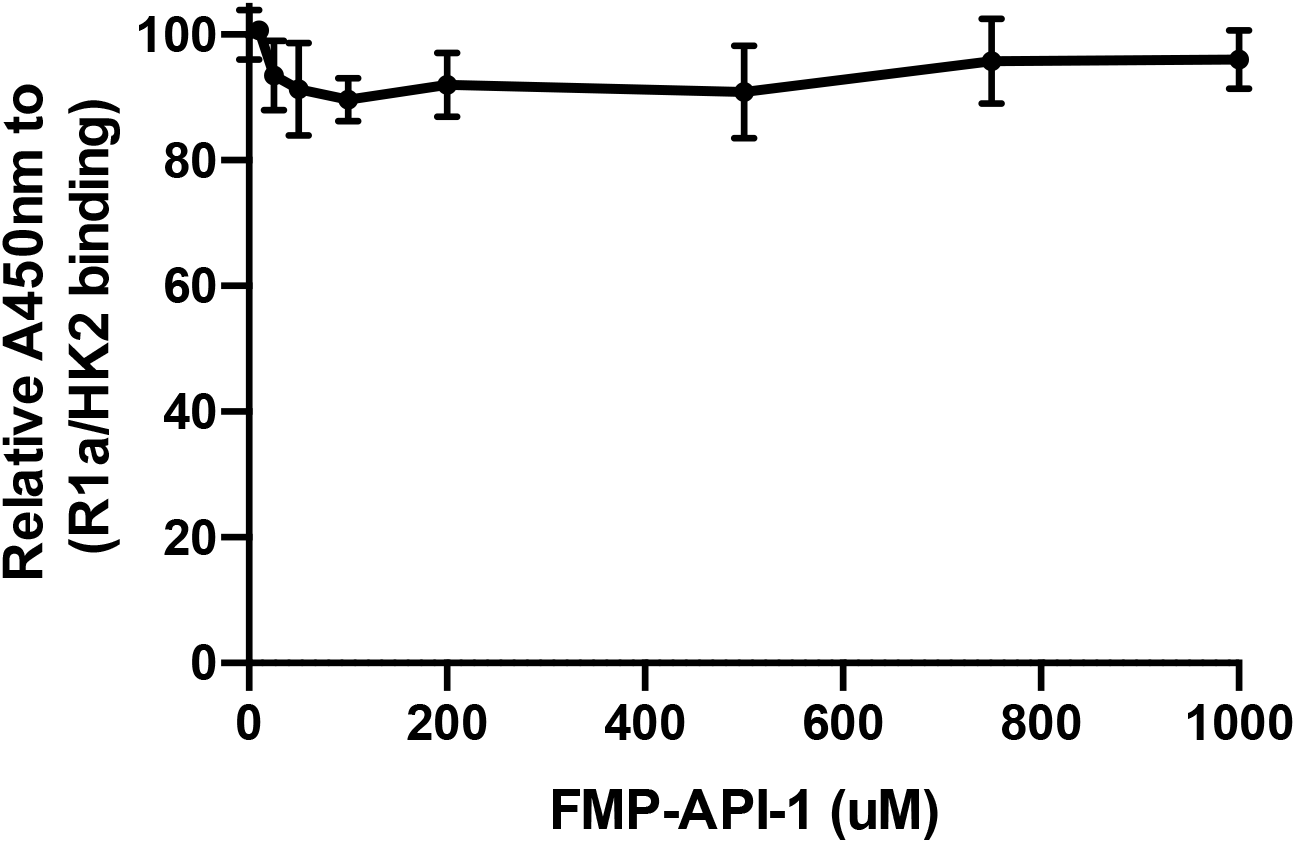
The effect of FMP-API-1 on HK2-PRKAR1a binding. Nickel-coated 96-well plates were incubated overnight at 4°C with His-PRKAR1α (50 nM) and then incubated with Myc-HK2 (32 nM) for 2h. Thirty minutes before the end of the later incubation, increasing concentrations of FMP-API-1 (0 – 1000uM) are added to the wells. HK2 binding was detected with anti-Myc-HRP conjugated antibody, and an HRP-catalyzed reaction with a chromogenic substrate solution.

**Extended Data Figure 8:**
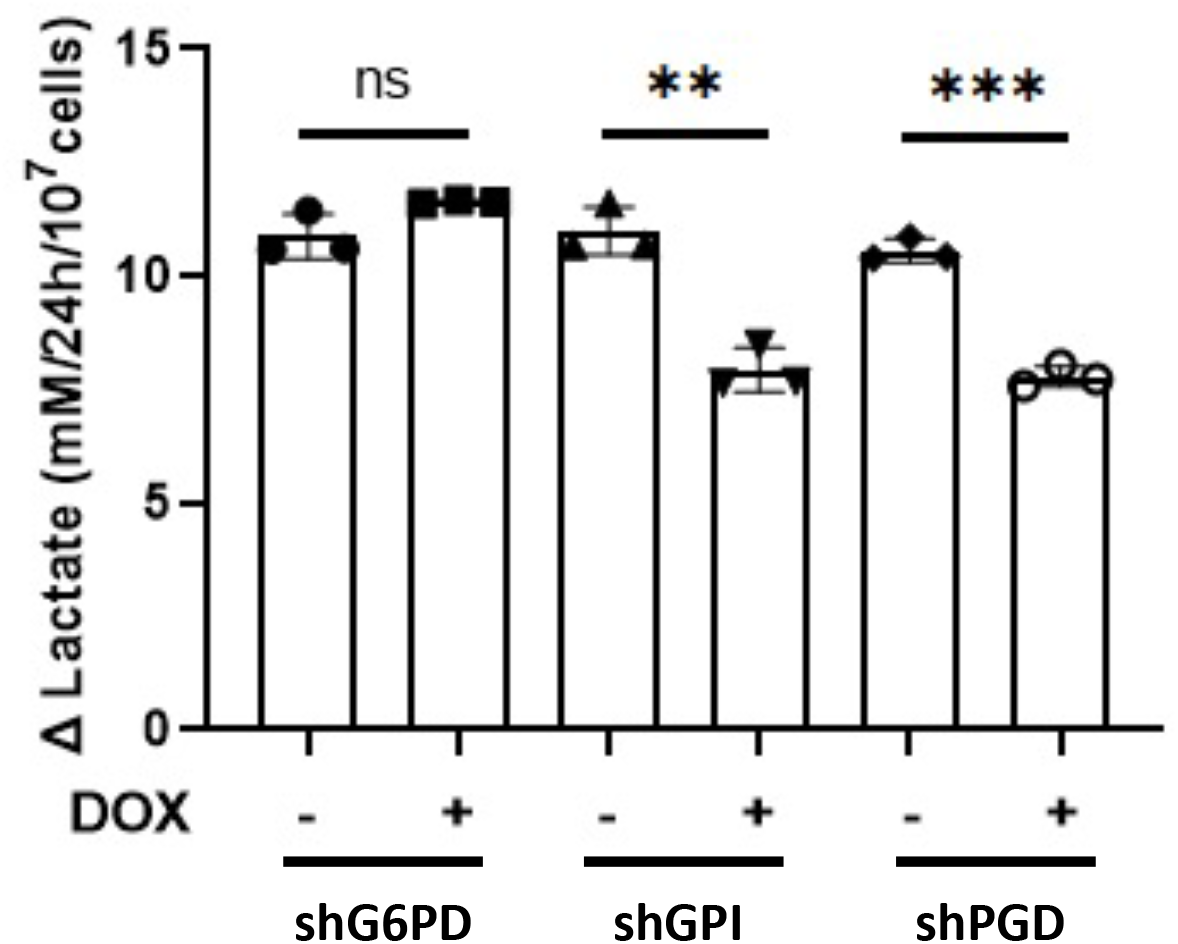
Extracellular lactate production rates in A549 cells. The amount of extracellular lactate production in 24 h was normalized to viable cell number; cells were treated with or without 0.2ug/ml DOX for 6 days. Results are the mean ± SEM of 3 independent experiments. **P<0.005, ***P<0.0005

**Extended Data Figure 9:**
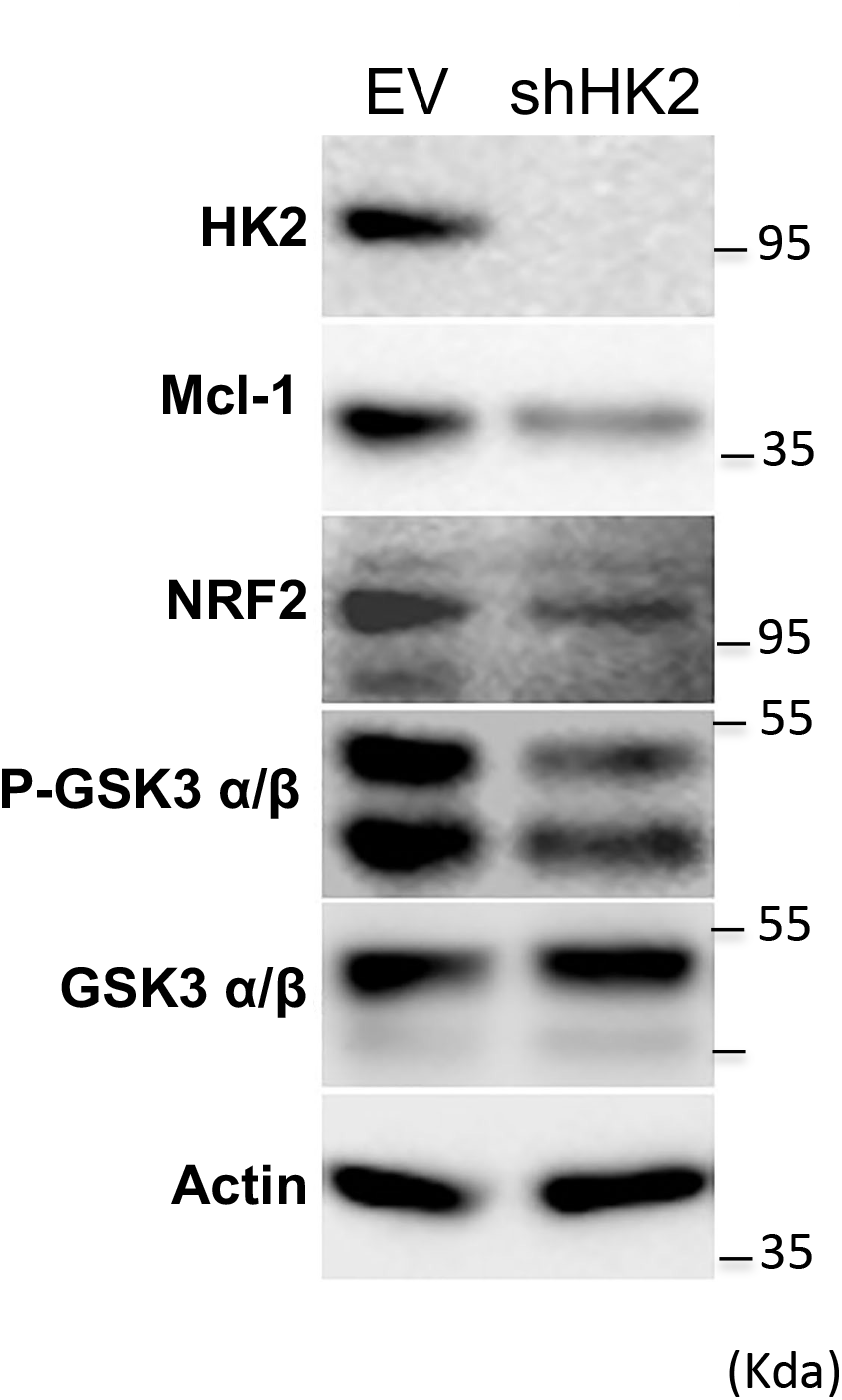
Representative immunoblot image showing P-GSK3β, Mcl-1 and NRF2 levels after the knockdown of HK2 in A549 cells.

**Extended Data Figure 10:**
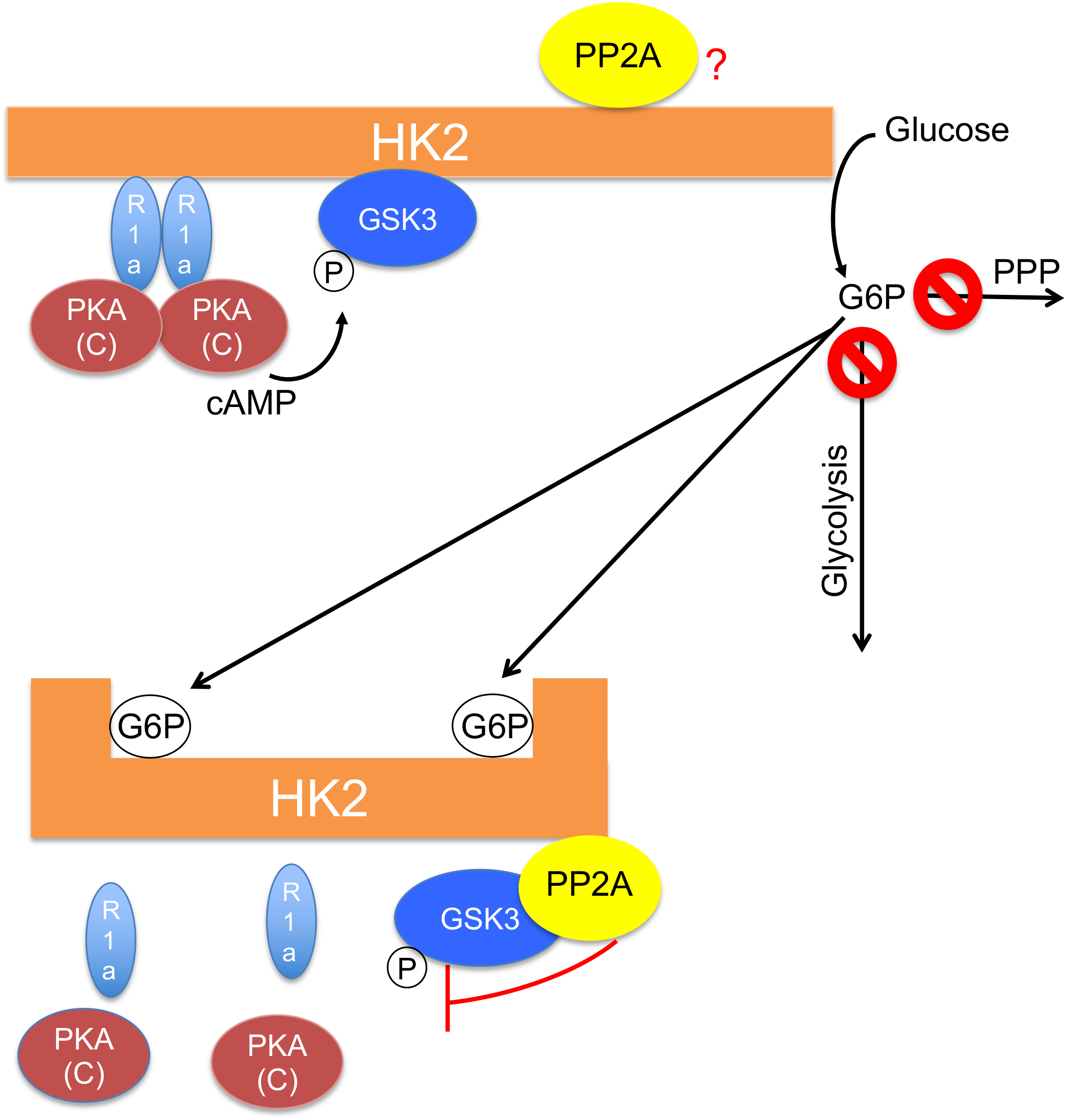
A model depicting HK2 as a scaffold for GSK3 and PRKAR1a-PKA. When cells have high glucose flux, HK2 brings PKA and GSK3 into close proximity. In the presence of cAMP, PKA is released from PRKAR1a to phosphorylate GSK3. When glucose flux is attenuated and G6P accumulates, an allosteric change is conferred to HK2 releasing GSK3 and PRKAR1a and increasing the availability of phosphorylated GSK3 to PP2A. It is possible that the amino-terminus half and the carboxy-terminus half of HK2 each binds GSK3 and PRKAR1a.

**Extended Data Figure 11:**
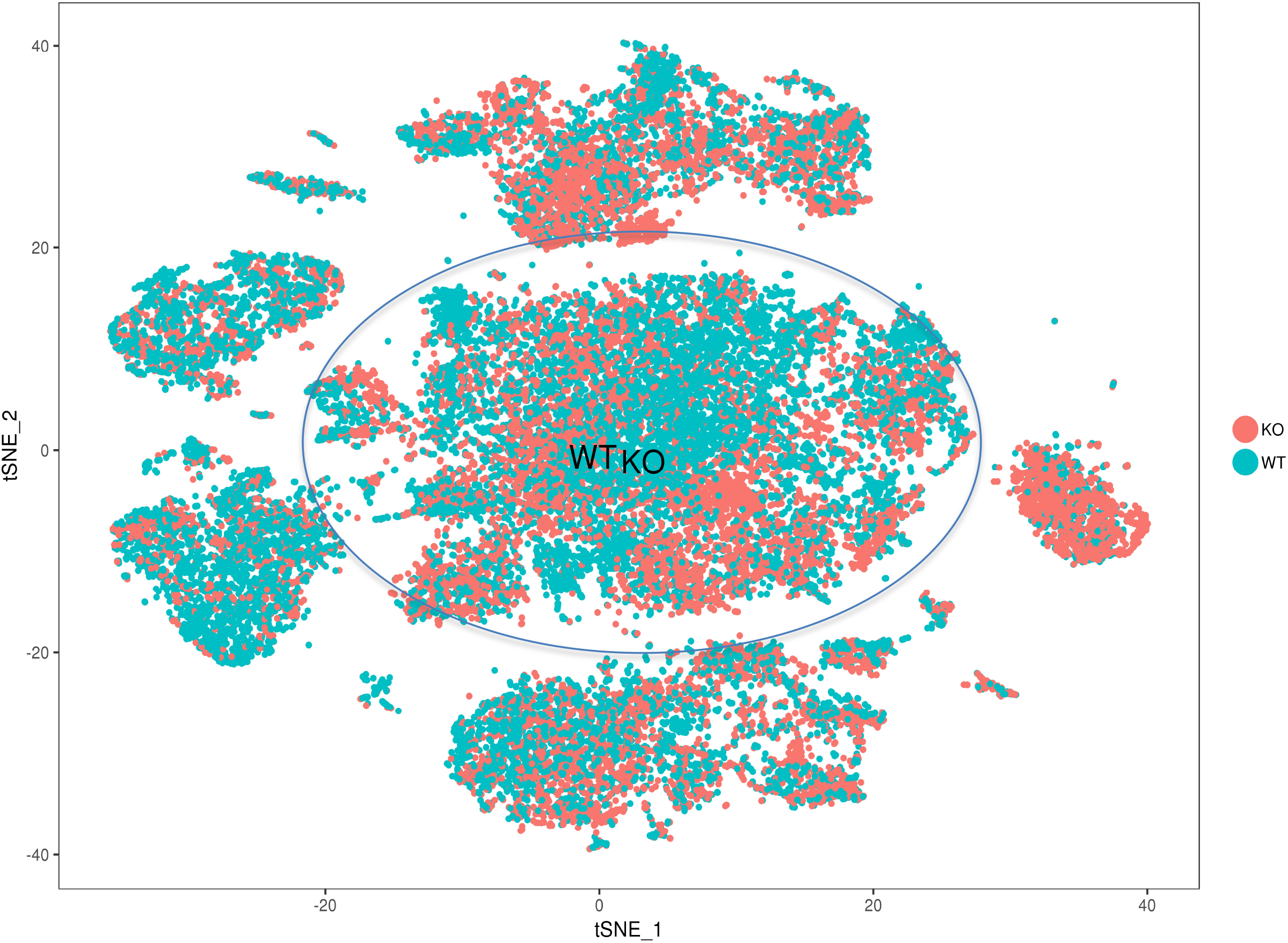
tSNE plot of pooled wild-type and HK2 deletion primary tumor samples. t-Distributed Stochastic Neighbor Embedding (tSNE) plot of pooled *MMTV-PyMT; HK2^f/f^* and *MMTV-PyMT; HK2^f/f^; UBC- Cre^ERT2^* primary mammary gland tumor samples (n=7 and n=8 respectively). There was no significant difference in the percentage of cells grouped into the clusters between HK2-deleted primary tumors (KO, red) and control primary tumors (WT, blue). Primary PyMT positive tumor cells are circled.

**Extended Data Figure 12:**
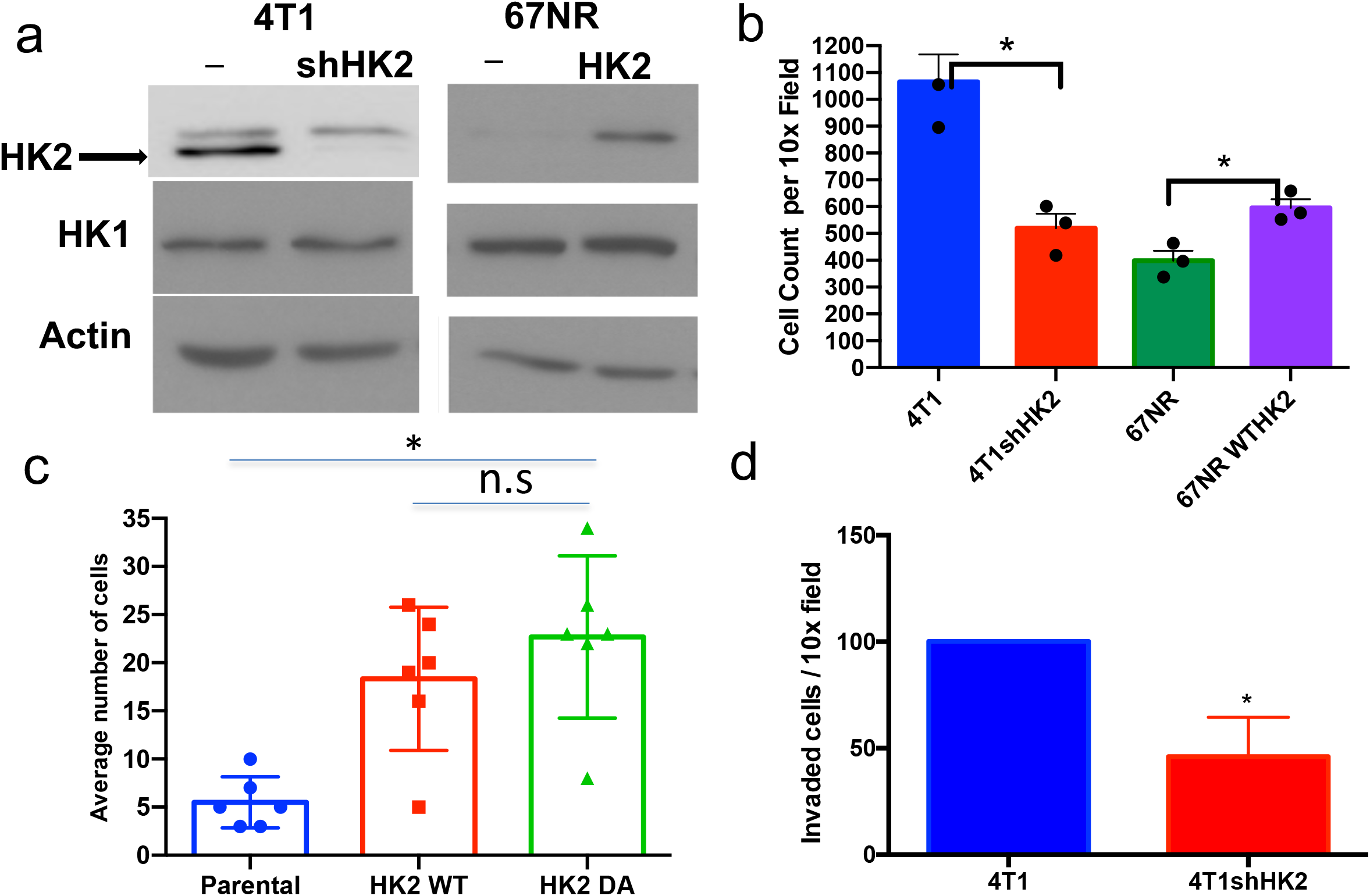
HK2 silencing in 4T1 cells decreases transwell migration and invasion, while overexpression of HK2 in 67NR cells increases transwell migration. **a.** Immunoblot showing HK2 levels in 4T1 cells before and after silencing of HK2 and HK2 levels 67NR cells before and after overexpression of HK2. **b.** Transwell migration comparing 4T1 shHK2 and 67NR WTHK2 cells to their parental cell lines. For transwell migration analysis, the cells were incubated in the upper transwell chambers for 12 hr with no serum while 20% serum was added to the lower chambers as a stimulant. Migrated cells in five random fields were counted after crystal violet staining, and three independent experiments were statistically analyzed. Quantified data represented as the mean ± SEM, *p < 0.1 from three independent experiments with each group plated in triplicate. **c.** Transwell migration analysis comparing 67NR cells to cells expression Wt HK2 or HK2DA mutant. **d.** Transwell invasion of 4T1shHK2 compared to their parental cell line. Briefly, the cells were incubated in a gel-coated transwell chambers for 24 hr with no serum, and 20% serum was added to the lower chambers as a stimulant. The same numbers of cells were plated on control transwell chambers for migration. The percent areas of invaded and migrated cells stained with crystal violet in ten random fields were counted, and the percentage of invaded cells was calculated. Three independent experiments were performed. The data represent the mean ± SEM, *p < 0.05.

**Extended Data Figure 13:**
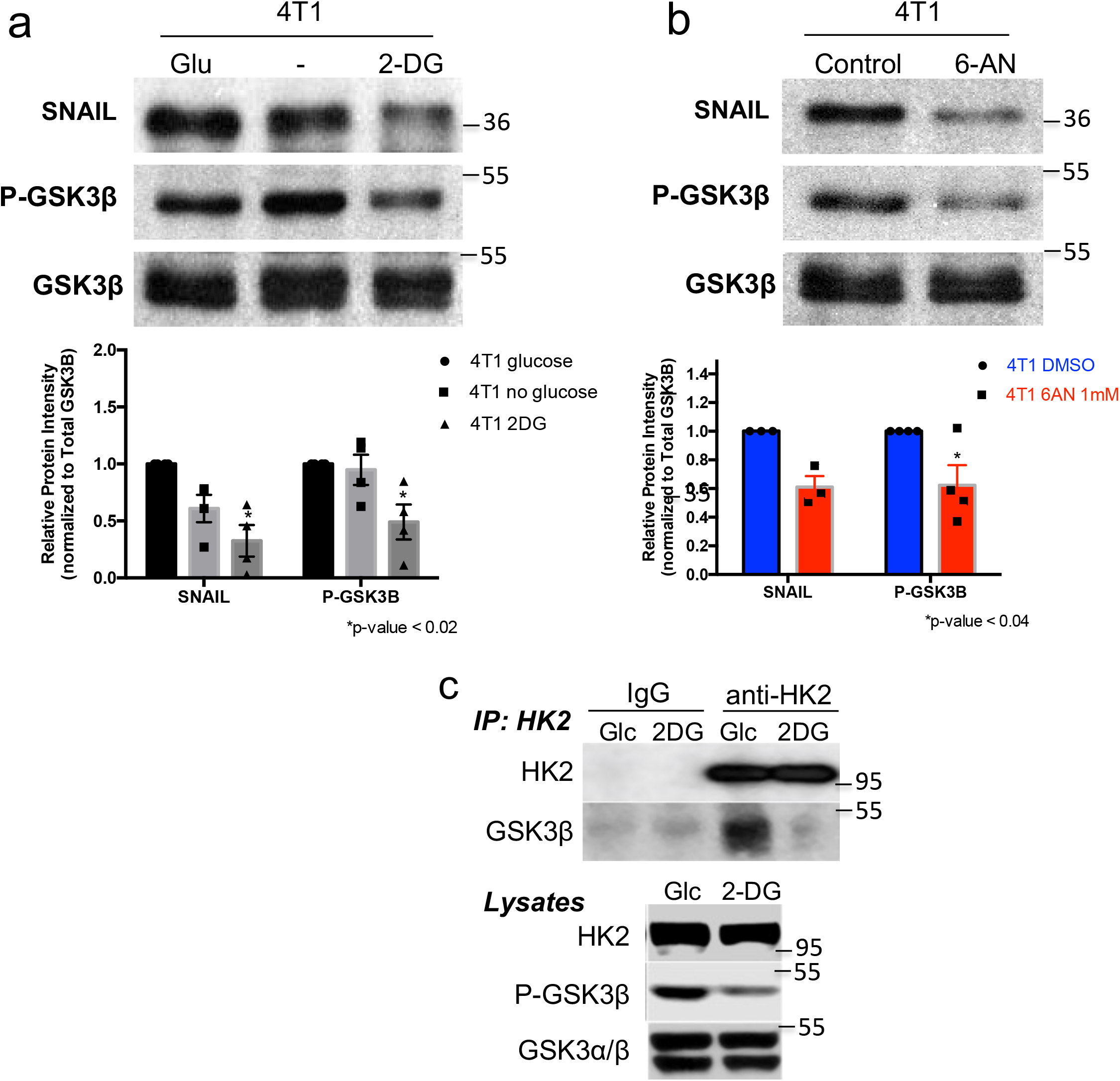
The effect 2-DG and 6AN treatment on SNAIL protein levels, GSK3 phosphorylation, and HK2- GSK3 binding. **a.** 4T1 cells were treated with 10 mM 2-DG or glucose as a control for 2 hr and then subjected to immunoblotting. Data were normalized to the amount of total GSK3β, and fold changes shown are relative to the control glucose-treated 4T1 cells. Quantified western blot data from 2-DG treatment are represented as the means ± SEMs. *p < 0.005 from triplicate experiments using an unpaired t-test (n=4). **b.** 4T1 cells were treated with 1 mM 6-AN for 12 hr and then subjected to immunoblotting to determine SNAIL and P-GSK3β levels. Quantified western blot data after 6-AN treatment are represented as the means ± SEMs. p < 0.03 from triplicate experiments using an unpaired t-test (n=4). Data were normalized to the amount of total GSK3β, and the fold changes shown are relative to the control DMSO-treated 4T1 cells. **c.** 4T1 cells were incubated in glucose free medium in the presence of 10mM glucose (Glc) or 2- DG. After 2hr, cells were lysed for immunoprecipitation. Endogenous HK2 was immunoprecipitated with anti-HK2 and subjected to immunoblotting with anti-HK2 and anti- GSK3β antibodies.

